# Critical events of the origins of life

**DOI:** 10.1101/2020.09.09.289678

**Authors:** Caleb Deen Bastian, Herschel Rabitz

## Abstract

We discuss some critical events of the origins of life using a mathematical model and simulation studies. We find that for a replicating population of RNA molecules participating in template-directed polymerization, the hitting and establishment of a high-fidelity replicator depends critically on the polymerase fitness and sequence specificity landscapes and on genome dimension. Probability of hitting is dominated by polymerase landscape curvature, whereas hitting time is dominated by genome dimension. Surface chemistries, compartmentalization, and decay increase hitting times. These results suggest replication to be the first ‘privileged function’ marking the start of Darwinian evolution, possibly in conjunction with clay minerals or preceded by metabolism, whose dynamics evolved mostly during the final period of the search.

## 1 Introduction

The origins of life (OOL), abiogenesis, is a matter of high importance, for it gives insight into the distribution of life in the universe. Most OOL models are based on ‘privileged functions’ that prioritize one of replication, metabolism, or compartmentalization (Lanier and Williams, 2017). In this vein, we focus on the RNA world hypothesis, where life begins with self-replicating RNA molecules that can evolve under Darwinian evolution (Orgel, 1968; Gilbert, 1986; Joyce, 2002). This is an information-centric perspective on abiogenesis which we refer to as ‘RNA-OOL,’ representing the putative beginning of Darwinian evolution. Other theories of abiogenesis prioritize metabolism (Dyson, 1999), cellular compartmentalization (Sakuma and Imai, 2015; Szostak et al., 2001), or hydrothermal vent chemical gradient energy (Martin et al., 2008). Information centrism interprets a living organism as an operating genetic communication system (GCS) in some connected domain that encodes and decodes genomic state relative to a replication channel.

While genomics, epigenomics, and transcriptomics of modern-day organisms are based on DNA, RNA, and epigenetic marks such as DNA methylation, RNA-OOL in its purest form concerns the dual-function of RNA as an informational polymer and ribozyme. Clues to the RNA world, among others, are found in the nucleotide moieties in acetyl coenzyme A and vitamin B12, the structure of the ribosome as a ribozyme (Cech, 2000) and moreover the centrality of RNA to the translation system, and the existence of viroids (Diener, 1971). The putative canonical RNA-OOL sequence involves RNA dependent RNA polymerase (RNAP) and Hammerhead (HH) ribozymes. It is important to note that this setup is unnecessary, as the first role of RNA may not have been template-assisted polymerization but instead based on mechanisms of RNA recombination and networking (Vaidya et al., 2012). If we assume the RNAP-HH genome is size 200 nucleotides, then there are 4^200^ ≈ 10^120^ possible sequences in this space of genomes. Starting from some population of interacting RNA molecules, we are interested in the hitting times of critical events. Evolution seems to have concluded this search for a high-fidelity replicator in a fairly short period of time, i.e. within 400 million years of the Earth having a stable hydrosphere (Joyce, 2002).

A subtlety to the RNA-OOL argument is that template-directed polymerization by its very nature essentially requires two copies of the genome, one for the polymerase and another for the template. This makes pure RNA-OOL improbable, requiring two independent hittings. An escape hatch is the fact that the clay mineral montmorillonite, which is common on Earth, has polymerase activity for RNA (Ertem and Ferris, 1997), so minerals furnish a genome complementary to the first, enabling template-directed polymerization to proceed. Hence we assume RNA-OOL takes place in an environment having clay. Some have proposed that clay not only promotes OOL but actually constitutes it, with the first living thing clay crystals, which then later gave rise to RNA-based replication (Cairns-Smith, 1987); however, such mineral life as a GCS has high noise.

Pre-RNA worlds have been suggested, with RNA being preceded or augmented by alternative informational polymers, such as other nucleic acids (Orgel, 2000), beta amyloid (Maury, 2015), polycyclic aromatic hydrocarbons (Ehrenfreund et al., 2006), lipids (Segré et al., 2001), peptides (Kunin, 2000), and so on. Note that many of these are exposed to the so-called code conversion problem, whereby a mapping must be established from one informational alphabet to another to ensure continuity of information of the genome. RNA and DNA alphabets need only a single substitution to convert between each other (along with ribose and deoxyribose) and hence the transition from RNA to DNA in information storage is simple in principle. Furthermore it is possible that pre-RNA worlds existed independent of the RNA world in the sense they are not ancestral to the RNA world, and that these worlds may have had non-trivial interactions with the RNA world. Thus are the possibilities.

NASA’s perspective on abiogenesis requires the following to be identified: abiotic sources of organic compounds, mechanisms of synthesis and function of macromolecules, energy sources, and environmental factors (Hays, 2015). Towards this, we assume the molecular inventories are provided by meteorites (KVENVOLDEN et al., 1970) or by Miller-Urey processes (Miller and Urey, 1959). For energy and environmental factors, we consider a variant of Darwin’s “warm little pond,” where the putative environment for origins of life is an icy pond with geothermal activity or perhaps a hydrothermal vent: ice and cold temperature facilitate complexing of single strands into double strands and polymerization (Attwater et al., 2013), and heat (energy) facilitates dissociation of double strands into single strands.

RNA-OOL has attracted many modeling efforts and analyses (Coveney et al., 2012; Lanier and Williams, 2017). Two-dimensional spatial modeling has been applied in which reactions occur locally with finite diffusion, suggesting a spatially localized stochastic transition (Wu and Higgs, 2012), simulated using Gillespie’s stochastic simulation algorithm (SSA) (Gillespie, 1977). Another model is of autocatalytic sets of collectively reproducing molecules, which has been developed in reflexively autocatalytic food-generated (RAF) theory (Hordijk and Steel, 2004). Various physics-based analyses have been conducted, such as in light of Bayesian probability, thermodynamics, and critical phenomenon (Walker, 2017). Systems of quasi-species based on the principle of natural self-organization called hypercycles employ non-equilibrium auto-catalytic reactions (Eigen and Schuster, 1977). Non-linear kinetic models for polymerization have been used to study the emergence of self-sustaining sets of RNA molecules from monomeric nucleotides (Wattis and Coveney, 1999).

We develop a simple mathematical model at the genome level to represent the synthesis and function of RNA molecules in order to gain insight into the hitting times of various critical events of the origins of life. The idea is to study the surface of hitting times in terms of the structure of the system. Let *H* = {Adenine, Uracil, Guanine, Cytosine}, abbreviated as *H* = {*A, U, G, C*}, corresponding to the RNA bases. We study the space of genomes having length *n, E* = *H*^*n*^ with *ℰ* = 2^*E*^, so (*E, ℰ*) is the measure space of all RNA sequences of length *n* with |*E*| = 4^*n*^. Appendix A gives overview of (*E, ℰ, v*). A related space, though not utilized in this article, is the space of all RNA sequences up to length *n*, 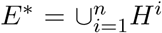 with ℰ* = 2^*E*\*^ and size 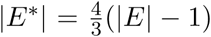. The model lacks many features of realism, such as genome size variability, finite sources of “food” (activated nucleotides in our context), limited diffusion rates, poor system mixing, chirality, and so on, in order to concentrate on the process as a search problem. A key concept is the notion of fitness and similarity surfaces or landscapes on (*E, ℰ*). The notion of such landscapes has been studied extensively in evolutionary biology (Wright, 1984). Landscape topology has been considered in an Opti-Evo theory, which assumes sufficient environmental resources and argues that fitness landscapes do not contain “traps” and globally optimal genomes form a connected level-set (Feng et al., 2012).

We describe the model in Section 2, where we define a replicating reaction network, whose random realizations are constructed using SSA. We describe hitting times as key random variables of interest and characterize polymerization as a transition kernel. In Section 3 we conduct and discuss simulation studies based on SSA, where we analyze the structure of the polymerase measures and the input-output and survival behaviors of the hitting times given the parameters of the system. In Section 4 we end with conclusions.

## 2 Materials and methods

We build a simplified mathematical model for the time-evolution of a population of interacting RNA molecules in solution. Let *X*_*t*_ be the population at time *t* ∈ ℝ_+_ with initial population *X*_0_. *X*_*t*_ is a multiset. We assume the system is well mixed and has access to an infinite source of activated nucleotides.

The complement of *x* ∈ *E* is denoted *x*^*c*^ ∈ *E*, attained using the base-pairing *A* with *U* and *C* with *G*. Let

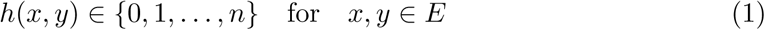

be the Hamming distance between *x, y* ∈ *E* as the number of positions in *x* and *y* where the nucleotides differ. We have *h*(*x, x*) = 0 and *h*(*x, x*^*c*^) = *n*.

### 2.1 Core model

We describe the reaction network of the system below.

#### System

We model the population of genomes which can form double-stranded helices, dissociate, and noisily replicate with replicator fitness and sequence specificity. We interpret each genome as a set, either containing one element—a single-stranded genome—indicated as {*x*}—or two elements, single and complementary stranded genomes, indicated as {*x*} ∪ {*x*^*c*^} = {*x, x*^*c*^} where *x, x*^*c*^ ∈ *E*. We define system elements as sets

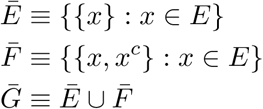

with respective *σ*-algebras, 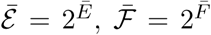 and 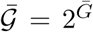. The reaction network of the system is given by

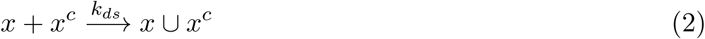

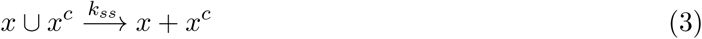

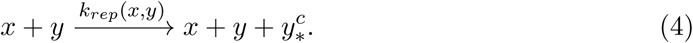

and expressed in terms of sets

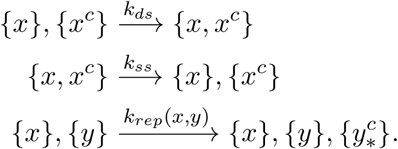

Reaction (2) is double-strand formation from complementary single-strands. Reaction (3) is the dissociation of double-strands into single-strands, caused by a heat source. Reaction (4) is template-directed polymerization of a single-strand (the template) by another single-strand (the polymerase), producing a single-strand complementary to the template with some fidelity.

We take

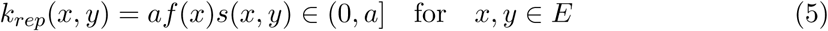

where *a >* 0 is a positive constant, *f* : *E* ↦ (0, 1] is the replicative *fitness* of *x* (as a polymerase) and *s* : *E* × *E* ↦ (0, 1] is the *similarity* between *x* and *y*, a symmetric function. The idea is that the replicator has fitness and specificity for sequences. Finally *x* replicates *y*, outputting a noisy version 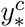, where each nucleotide position has fidelity probability *p* : *E* ↦ (0, 1]. This is a continuous-time branching model with mutation. We simulate trajectories of the system using SSA (Gillespie, 1977).

The system is assumed to be well-mixed, and all parts of the system are accessible for reactions. This means that reactions across various disjoint volume elements of the system are dependent. We do not consider the effects of finite diffusion, which effects a length scale above which disjoint volume elements are effectively independent (Wu and Higgs, 2012). Basically limited diffusion breaks the system into non-interacting regions, whereby reactions are localized.

#### High-fidelity set

Pick an arbitrary subset of genomes *R* ⊂ *E* as high-fidelity replicators of size *r* = |*R*|. We define *R* two ways.

We define *R* using a product of non-empty random nucleotide subsets {*A*_*i*_ ⊆ *H* : *i* = 1,…, *n*} for each nucleotide position

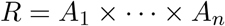

so that

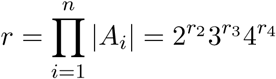

where *r*_2_ = |{*A*_*i*_ : |*A*_*i*_| = 2}|, etc. and *r*_1_ + *r*_2_ + *r*_3_ + *r*_4_ = *n*. For simplicity, we assume |*A*_*i*_| ∈ {1, 4} with fraction 4 being *q* ∈ (0, 1). Thus *R* is a subset of *E* defined as a product space. This model is essentially single-nucleotide polymorphism (SNP), where each SNP position takes values of any of the nucleotides.

For another construction of *R*, we define a finite union of *m* random genomes *R* = {*x*_1_,…, *x*_*m*_}.

#### Distance

Define the Hamming distance *H* between *x* and *R* as

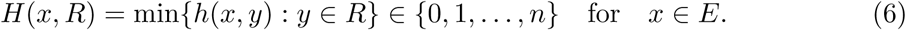

#### Fitness

Define “tent-pole” fitness *f*_*k*_ of *x* ∈ *E* and *R* for *k* ∈ ℝ_+_ as

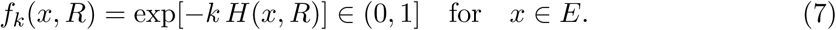

The maximums are the genomes of *R*, which are the “points” or “poles” of the surface, with exponential decay into the remainder of the space in string distance. The strength of the decay is governed by parameter *k*, which can be specified through the value of fitness at *H*(*x, R*) = *n* (genome dimension),

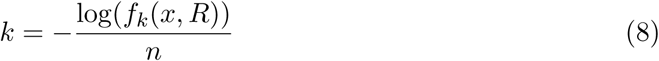

Appendix B describes other fitness functions.

#### Similarity

Symmetric distance *S* is defined as

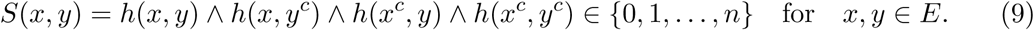

Similarity *s*_*k*_ of *x, y* ∈ *E* for *k* ∈ ℝ_+_ is defined in terms of exponential decay as

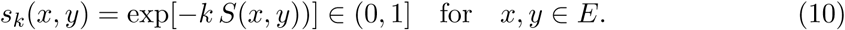

Similarity is a measure of the distance between the replicator and the template or template complement, where smaller distances convey higher similarities. Various parameterizations of similarity may be considered.

#### Fidelity

Finally, polymerization fidelity probability for *k* ∈ ℝ_+_ is defined as

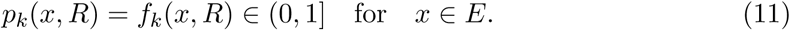

Note that *f*_*k*_(*x, R*) = *p*_*k*_(*x, R*) = 1 for *x* ∈ *R*.

Note that fitness, similarity, and fidelity are defined for single-stranded genomes (*E, ℰ*).

#### Counting representation

The process 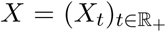 is the time-evolution of the system. *X*_*t*_ contains the individual single stranded molecules in the set *Ē* and double stranded molecules in the set 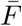, with set 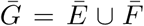 having size 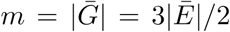. Note that {*x, x*^*c*^} = {*x*^*c*^, *x*}, so 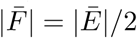. *X*_*t*_ induces a random counting measure *N*_*t*_ on 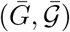 as

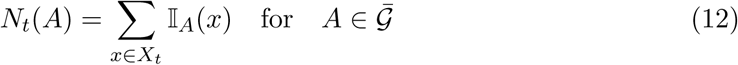

The total count is 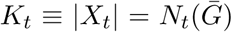. We assume that the counter *N*_*t*_ is maintained for all *t* ∈ ℝ_+_.

#### Reaction rates

The total reaction rate is given by

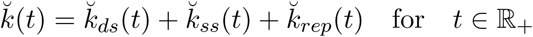

where

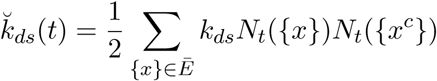

and

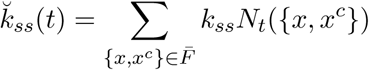

and

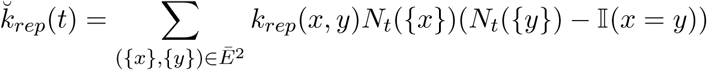

The first reaction rate is a sum over *Ē* with size 4^*n*^. The second a sum over 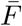 with size 4^*n*^*/*2. The third reaction is a sum over *Ē*^2^ with 4^2*n*^ = 16^*n*^ elements. Therefore 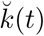 requires 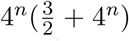 elements to be evaluated for every reaction. Clearly direct representation on the full space is very expensive and impractical for even modest *n*. One obvious way to improve efficiency is not summing over the zero elements. Form sets

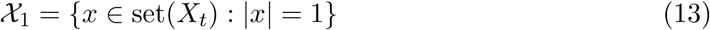

and

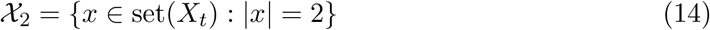

Then direct calculations are

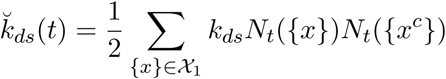

and

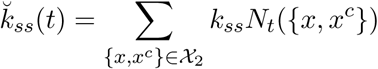

and

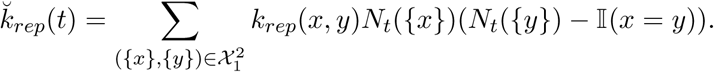

For this approach the replication rate has quadratic dependence on |*𝒳*_1_|. Using the reaction rates, the system may be exactly simulated using SSA. The reaction at time *t* with rate 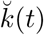 occurs over time interval 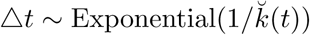. As 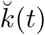 increases with increasing 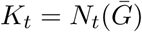, the reaction rate increases and reaction duration Δ*t* decreases over time. The natural consequence of increasing process intensity is that the system speeds up.

The quadratic dependence may still be too expensive for large simulations. Appendix D describes Monte Carlo approximation of the reaction rates.

### 2.2 Hitting times

We define some hitting times. The initial population consists of *I* single-stranded genomes *X*_0_. Define hitting time *τ* for the time of the first replication event

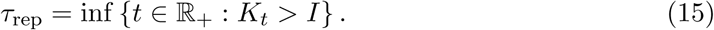

Define the hitting time *τ* for the appearance of *R*,

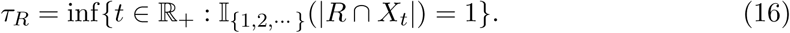

Put *X*_*R*_ = *R* ∪ {{*x, x*^*c*^} : *x* ∈ *R*} and define the volume fraction of *R* at time *t* as

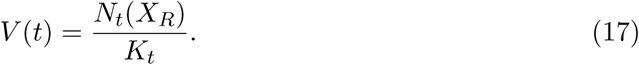

Define hitting time *τ* where *R* emerges and reaches a minimum volume fraction,

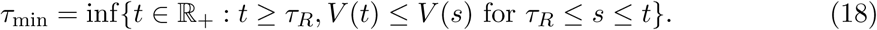

Define hitting time *τ* for the time *R* constitutes some volume fraction *v* ∈ (0, 1] of the population,

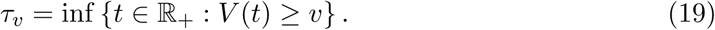

In practice for simulations, *τ*_*v*_ is censored based on some total number of reactions, that is, if the volume fraction is not achieved by *n* reactions, *τ*_*v*_ = *∞* because there is no arrival time.

We have that *τ*_rep_ *≤ τ*_*R*_ *≤ τ*_min_ *≤ τ*_*v*_.

For SSA, we specify a maximum number of reactions *N* to simulate. We have parameters *θ* ∈ Θ for *τ*, such as landscape curvature *k*, genome dimension *n*, etc. Therefore *τ* (*θ*) is right-censored with value *∞* at simulation time *a*, as some simulations will stop at time *a* with no arrival time. These are censoring events. For fixed *θ* the *τ* (*θ*) is a random variable, due to the stochastic nature of SSA. Hence for each *θ*, we attain a set of *M* realizations of *τ* as

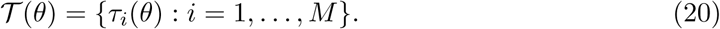

For convenience we assume the *𝒯* (*θ*) is ordered by non-censored followed by censored.

#### Functional structure

For each *θ*, we record two values: the number of hitting events in *𝒯*

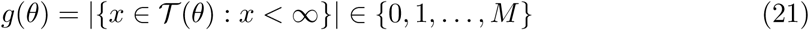

and the average of *τ*

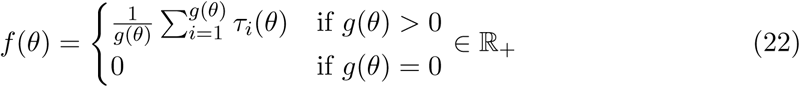

To describe the functional structure of *f* (*θ*), we require a classifier which determines whether or not *g*(*θ*) = 0 and a regressor for the value of *f* (*θ*) for *g*(*θ*) *>* 0. Assume that the parameters *θ* = (*θ*_1_, …, *θ*_*n*_) are randomly sampled according to distribution *v* = ∏_*i*_ *v*_*i*_ and the hitting times recorded. High dimensional model representation (HDMR) may be attained for the classifier (as a probabilistic discriminative model) and the regressor of *f* (*θ*). For the regressor, we have HDMR

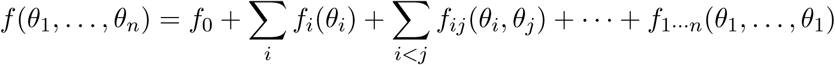

The HDMR component functions {*f*_*u*_} convey a global sensitivity analysis, where defining variance term

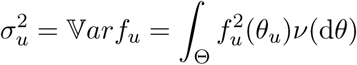

we have a decomposition of variance

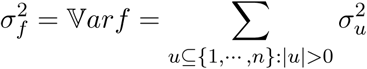

The normalized terms 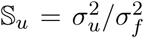 are called sensitivity indices. Appendix E gives brief description of global sensitivity analysis via HDMR.

#### Statistical structure

A second analysis can be conducted on *𝒯* (*θ*) for the *θ* using reliability theory. Put ℐ(Θ) *≡* {*𝒯* (*θ*) : *θ* ∈ Θ} where Θ = {*θ*_*i*_} is an independency of parameter values. For each *θ* ∈ Θ we partition *𝒯* (*θ*) into *C* censored values with censor times *𝒞*(*θ*) = {*a*_*i*_(*θ*)} and *M−C* non-censored (hitting) values *𝒩* (*θ*) = {*x* ∈ *𝒯* (*θ*) : *x < ∞*}. The likelihood is given by

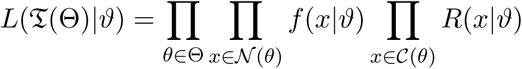

where *f* is the hitting time probability density function (‘failure density’) and *R* is censoring time distribution (‘reliability distribution’) and *ϑ* are the parameters of the density and distribution functions. Note that *f* and *R* each specify each other, so *ϑ* are the common parameters. Reliability definitions are given in Appendix F.

We show reliability quantities in Table 1 for the two-parameter (*α, β*) ∈ (0, *∞*)^2^ Weibull(*α, β*) distribution and Cox proportional hazard’s model where *γ* is a vector of coefficients for the *θ*. We use the Python software *lifelines* for estimation of *ϑ* for the Weibull-Cox model from data (Davidson-Pilon, 2019). The mean failure time 𝔼*τ* (*θ*) is given by

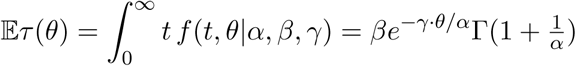

**Table 1:**
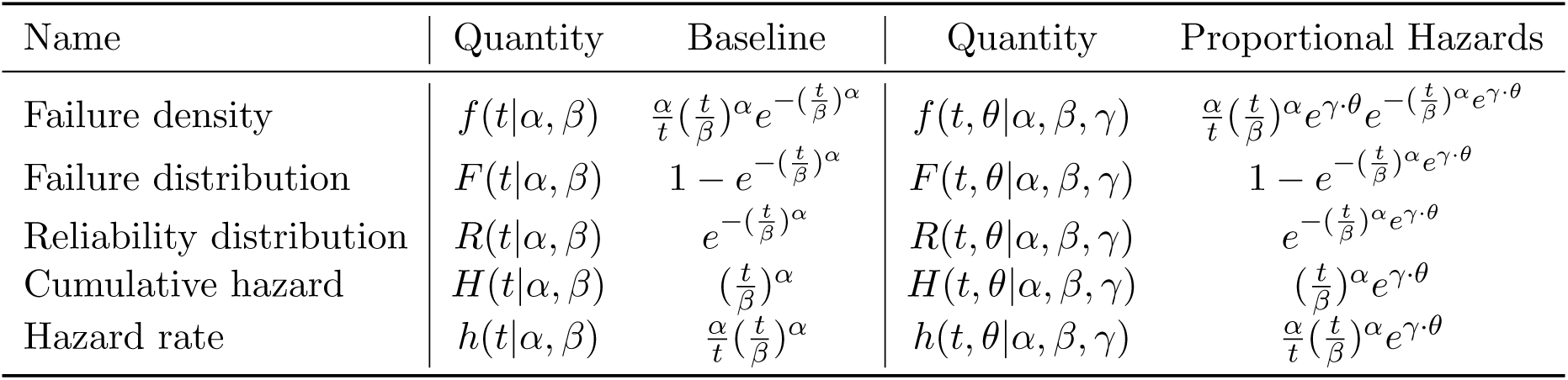
Weibull reliability model

We have second moment

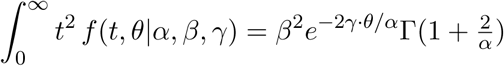

giving variance

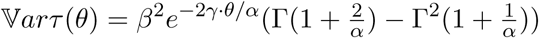

Thus if *γ <* 0, then 𝔼*τ* (*θ*) and 𝕍*arτ* (*θ*) exponentially increase in *θ*.

### 2.3 Hitting cardinality

We index the *N* reactions of *X*_*t*_ with arrival times {*T*_*i*_} in 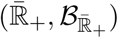 where 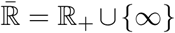. Define the hitting reaction *ϖ*(*θ*) ∈ ℕ in terms of hitting time 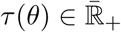 as

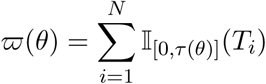

*ϖ*(*θ*) is right-censored at *N* reactions. If *τ* (*θ*) = *∞*, then *ϖ*(*θ*) = *N*. If *τ* (*θ*) *< ∞* then *ϖ*(*θ*) < ∞.

### 2.4 Surface chemistries

The system given by (2) of noisy polymerization requires two separate hitting events, one *x* ∈ *R* and either *x* ∈ *R* or *x*^*c*^ ∈ *E* in order for high-fidelity replicators to maximally engage in templated-directed polymerization and achieve some fraction of the population. This setup of RNA polymerase action, requiring two such events for the polymerase and template, makes the hitting times long. Basically the same information must be discovered twice before it can be used, which is unsatisfactory. Other molecules have RNA oligomerization and polymerase activities, such as the clay montmorillonite (Ferris, 2006; Ertem and Ferris, 1997), that are naturally found on Earth in various environments. Here the surface chemistry of montmorillonite performs oligomerization and polymerization of RNA molecules. The reactions are given by

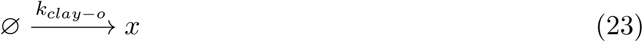

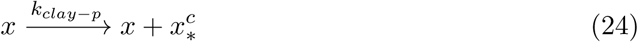

where polymerization is noisy with fidelity probability *p* ∈ (0, 1] and 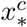 is the noisy complement of *x*. The reaction rates are given by

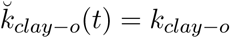

and

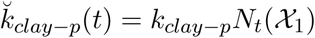

Therefore, upon the first hitting of *R* with genome *x* ∈ *R* through (4) or (23), *x* gives two high-similarity single-stranded genomes *x* and 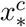 through (24), which then may participate in template-directed RNA polymerization (4).

### 2.5 Reactions as measure-kernel-functions

All the reactions *x* ↦ *y* which involve substrate *x* may be represented using transition kernels, which form linear operators. At each iteration of SSA, a reaction type is chosen, followed by a transition to a particular domain (*X, 𝒳*) with distribution *v*_*t*_, followed by mapping into a codomain (*Y, 𝒴*) using transition probability kernel *Q* with distribution *µ*_*t*_ = *v*_*t*_*Q*. The notions of *v*_*t*_ and *Q* involve *measure-kernel-functions*. The probability of transition of *y* into *B* ∈ *𝒴* given *x* is given by *Q*

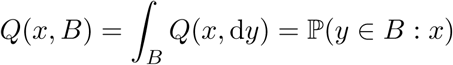

Appendix C recalls some facts about *Q*.

We index the reaction types on (*Z, 𝒵*) = (ℕ_*>*0_, 2^ℕ^*>*0). Let *η*_*t*_ be the probability measure on (*Z, 𝒵*) formed from the normalized reaction rates. Let *Q*_*\**_ be the transition kernel from (*Z, 𝒵*) into (*X, 𝒳*). Then *η*_*t*_*Q*_*\**_ = *v*_*t*_ is the distribution on (*X, 𝒳*) and *µ*_*t*_ = *v*_*t*_*Q* is the distribution on (*Y, 𝒴*). Then a reaction is the mapping (*Z, 𝒵*) ↦ (*X, 𝒳*) *1*↦ (*Y, 𝒴*).

We define kernels *Q* for RNA and clay polymerization to provide insight into the reactions. Consider *X*_*t*_ for some *t* ∈ ℝ_+_. Recall *N*_*t*_ is the random counting measure of *X*_*t*_ on single and double-stranded genomes (*E* ∪ *F*, 2^*E*∪*F*^). For RNA and clay polymerization we take *v*_*t*_ as a (random) probability measure for *x* in domain (*X, 𝒳*) and describe a transition probability kernel *Q* from *x* into *y* in codomain (*Y, 𝒴*).

#### RNA polymerization

For RNA polymerization we have that

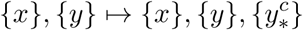

which results in the creation of the single-stranded genome 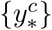. The first dimension is the polymerase and the second is the template.

**Definition 1** (Measure on domain *v*_*t*_). *Let v*_*t*_ *be a random probability measure on* (*E* × *E, ℰ* ⊗ *ℰ*) *formed by random counting measure N*_*t*_ (12)

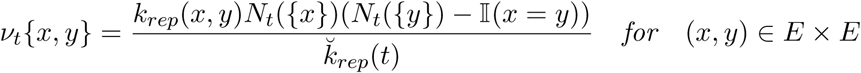

*with*

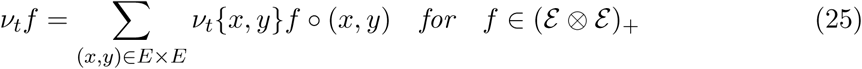

*We write v*_*t*_(*A*) = *v*_*t*_𝕀_*A*_ *for A* ∈ *ℰ* ⊗ *ℰ*.

Because the first two coordinates are preserved under the mapping, we focus on the new dimension as a transition from (*E* × *E, ℰ* ⊗ *ℰ*) into (*E, ℰ*) using transition probability kernel *Q*. In this case, *Q* is defined by a 16^*n*^ × 4 ^*n*^ matrix whose rows vectors (dimension 4 ^*n*^) are probability vectors. The structure of *Q* follows from the polymerase noisy replication, whereby each nucleotide position has fidelity probability *p* : *E* ↦ (0, 1], which depends on the first dimension of *E* × *E*. We put *p*_*x*_ = *p*(*x*) for *x* ∈ *E*.

##### Theorem 1

(Error distribution). *The number of errors by polymerase x* ∈ *E on template y* ∈ *E is distributed*

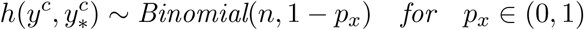

*with mean n*(1 *− p*_*x*_) *and variance np*_*x*_(1 *− p*_*x*_) *and*

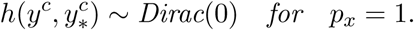

Partition *E* into level sets (*H*_*i*_(*y*)) by Hamming distance to the template complement

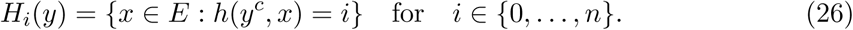

We define *Q* for RNA polymerization. *Q* completely encodes RNA polymerization using Theorem 1.

##### Corollary 1

(Transition probability kernel *Q*). *We have that*

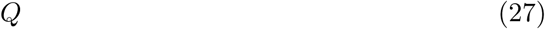

*for RNA polymerization is defined by*

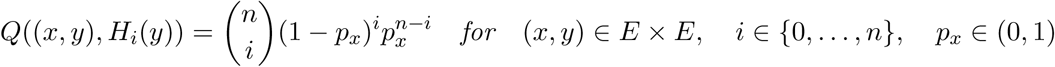

*and*

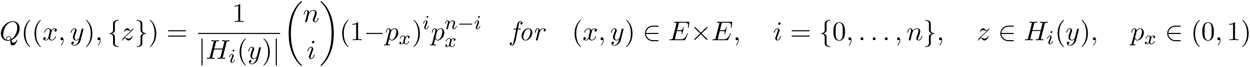

*and*

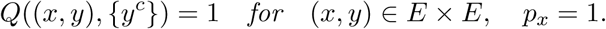

RNA polymerization is defined by *Q* using the binomial structure of polymerase error. A more sophisticated model could be defined as a sum of Bernoulli random variables with varying success probabilities in the Poisson binomial distribution. This could be used to take into account polymerase error that varies with nucleotide position, such as taking into account schemata such as repeats which destabilize the polymerase (Bornholt et al., 2016).

##### Proposition 1

(Measure on codomain *µ*_*t*_). *µ*_*t*_ = *v*_*t*_*Q is a probability measure on* (*E, ℰ*) *defined by*

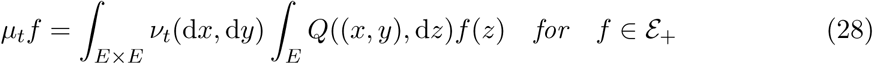

*It is multiplication of v*_*t*_ *as a* 16^*n*^ *dimension row vector with* 16^*n*^ × 4^*n*^ *dimension matrix Q, giving a* 4^*n*^ *dimension row vector v*_*t*_*Q. We write µ*_*t*_(*A*) = *µ*_*t*_𝕀_*A*_ *for A* ∈ *E*.

Define the partition (*H*_*i*_) of *E* as

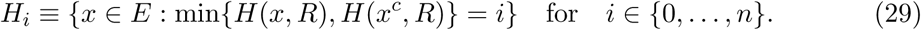

Then *µ*_*t*_ (*H*_*i*_) for *i* ∈ (0, …, *n*) is the distribution on genomes by distance to *R*, i.e.

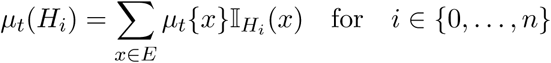

contains the instantaneous information of RNA polymerization.

**Clay polymerization** For clay polymerization, we have that

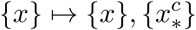

which we regard as a mapping from (*E, ℰ*) into (*E, ℰ*). Let *v*_*t*_ be a probability measure on (*E, ℰ*) defined by

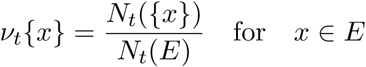

Similar to RNA polymerization, for clay fidelity probability *p* ∈ (0, 1] we have *Q* as the 4^*n*^ × 4^*n*^ matrix defined by

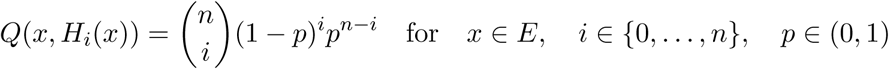

and

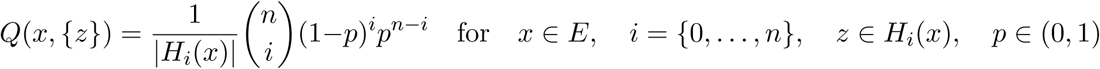

and

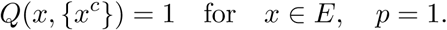

*µ*_*t*_ = *v*_*t*_*Q* is a probability measure on (*E, ℰ*) defined by

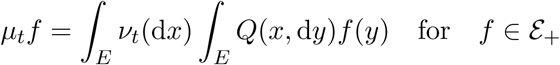

Note that, for SSA, *Q* is fixed over the simulation, whereas *v*_*t*_ depends on time. That is, the reactions are chosen according to the reaction rates, and the reactions each use respective *Q*. The *v*_*t*_ is formed using a random counting measure, so *v*_*t*_ is random. This approach generalizes in the obvious way to all the reactions.

### 2.6 Deterministic model

Consider the core model defined in (2), (3), and (4) with reaction rates 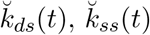, and 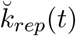. We are neglecting clay and decay for the moment. All the reactions impact (*E, ℰ*). For double-strand formation, the input space is (*X, 𝒳*) = (*E* × *E, ℰ* ⊗ *ℰ*) and the output space is (*Y, 𝒴*) = (*F, ℱ*). For double-strand dissociation, the input and output spaces are swapped. For polymerization, (*X, 𝒳*) = (*E*×, *ℰ* ⊗ *ℰ*) and (*Y, 𝒴*) = (*E, ℰ*). Hence (*E, ℰ*) is positively impacted by polymerization, positively impacted by double-strand dissociation, and negatively impacted by double-strand formation. Put [*x*] = *N*_*t*_({{*x*}}) and [*x, x*^*c*^] = *N*_*t*_({{*x, x*^*c*^}}). Recall the *v*_*t*_, *Q*, and *µ*_*t*_ = *v*_*t*_*Q* for the reactions, e.g., 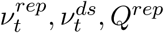, etc.

For *x* ∈ *E*, we have the system of 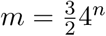 deterministic non-linear ordinary differential equations (ODEs) in *m* variables as mean-field equations

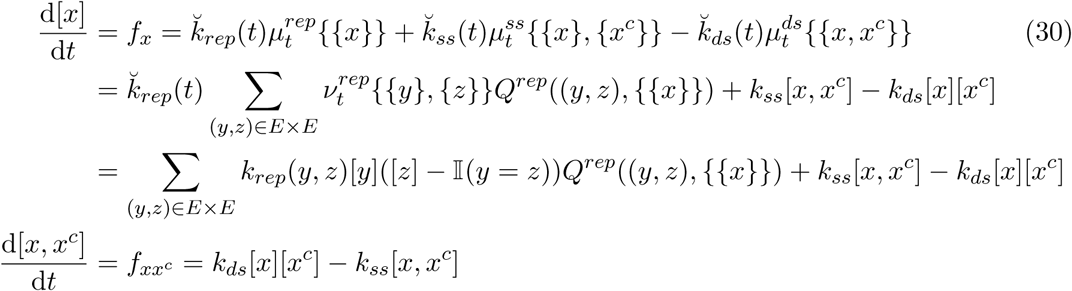

The fixed points of *f* are the equilibria of the system, i.e. *f* (*x*) = **0** for 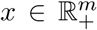. The Jacobian of the system is

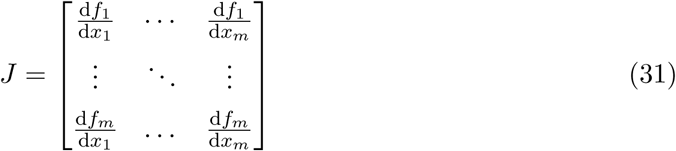

The eigenvalues of the Jacobian reveal the stability of the fixed points. If all the eigenvalues of the Jacobian evaluated at the fixed point have negative real part, then the fixed point is stable. If none of the eigenvalues are zero and at least one of the eigenvalues has positive real part, then the fixed point is unstable. If at least one eigenvalue is zero, then the fixed point can be either stable or unstable.

### 2.7 Decay

The RNA genomes have finite lifetimes in reality. This comes from a variety of sources, including radiation, pH, intrinsic molecular stability, etc. Assume double-stranded RNA is stable, whereas single-stranded RNA is not. Therefore we create a reaction for decay of single-stranded RNA into constitutive nucleotides

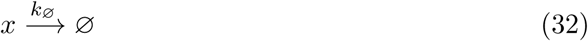

with reaction rate

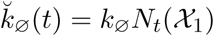

### 2.8 Compartmentalization

It is thought that compartmentalization plays a role in the origins of life, giving foci of reproducing genomes (Kitadai and Maruyama, 2018; Szostak et al., 2001). This is somewhat anticipated by the similarity measure *s*, where genomes are more likely to copy similar genomes than less similar ones. Explicit spatial effects may be modeled by assuming each *x* ∈ *E* is marked with a position on a bounded subset of the real line ([*−T, T*], *ℬ*_[*−T,T*]_) ⊂ (ℝ, *ℬ* _ℝ_). We think of this as a one-dimensional projection of the three-dimensional system. Additional species can be introduced, such as lipids, with reactions forming a micelle *M* (micellisation), which encloses some *A* = [*r, s*] ⊂ [*−T, T*]. We assume the lipids interact with the single stranded genomes in *A* to form micelles as

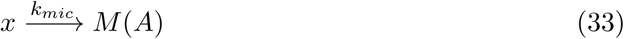

with reaction rate

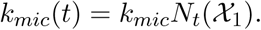

Hence micellisation is coupled to the population of genomes by design so that it evolves on roughly the same time-scale as genome activities. Note that micelles can enclose one another, i.e. *M* (*A*) and *M* (*B*) where *A* ⊂ *B* or *B* ⊂ *A*, but cannot cross, i.e. for all micelles at locations *A*,…, *B* we have that *A ∩ B* ∈ {*A, B*, ∅}. For example, suppose one micelle *A* = [0, 1] encloses another two, 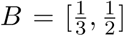 and 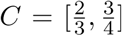. Then 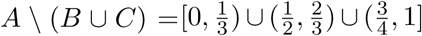. Although *A\* (*B* ∪ *C*) is disconnected in one dimension, the intervals are physically connected in three dimensions, where micelles are spheres. We identify each micelle to a union of disjoint intervals, disjoint across the micelles.

Let 𝔐 = {*A* ⊂ [*−T, T*]} be the collection of disjoint micelle regions. Let *l* : *G* ↦ [*−T, T*] be the location function of the genome. Let *L*_*t*_ = {*l*(*x*) : *x* ∈ *X*_*t*_} be the location set of the genomes. Then the random measure *N*_*t*_ on (*G, 𝒢*) is expanded to *M*_*t*_ on (*G* × [*−T, T*], *𝒢* ⊗ *ℬ*[*−T,T*]),

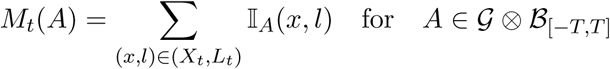

### 2.9 Metabolism

Metabolism is often promoted in origins-of-life (Dyson, 1999). For our purposes here, we shall identify metabolism reaction-state to the measurable space (*M, ℳ*) with probability measure *v*. Let *Q*_*m*_ be a transition kernel from (*M, ℳ*) into (*E, ℰ*), positing that metabolism precedes replication. For example, certain metabolic state may be precursors to the synthesis of RNA. Consider product space (*M* ×*E, ℳ*⊗ *ℰ*) with measure *µ* = *v*×*Q*_*m*_. Now we suppose that, upon achieving Darwinian evolution in replicators, the replicators will eventually become adapted to (*M, ℳ*). Hence we interpret (*M, ℳ*) as a mark-space of (*M* × *E, ℳ* ⊗ *ℰ*), representing genomic adaptation. Let *Q*_*e*_ be a transition kernel from (*M* × *E, ℳ* ⊗ *ℰ*) into (*M, ℳ*). Then *µ* = *v* × *Q*_*m*_ × *Q*_*e*_ is a probability measure on (*M* × *E* × *M, ℳ* ⊗ *ℰ* ⊗ *ℳ*), where

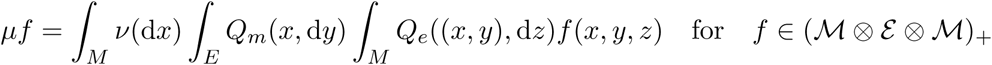

or

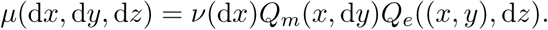

Therefore metabolism-first followed by replication and genomic adaptation is encoded by the structures of *Q*_*m*_ and *Q*_*e*_. We do not specify these transition kernels in this article but comment they are richly textured.

### 2.10 Reaction overview

The reactions of the system having decay and clay and their reaction orders are shown in Table 2. There is one zero-order reaction, three first-order reactions, and two second-order reactions. Additionally we show reactions and orders for compartmentalization and metabolism.

**Table 2:**
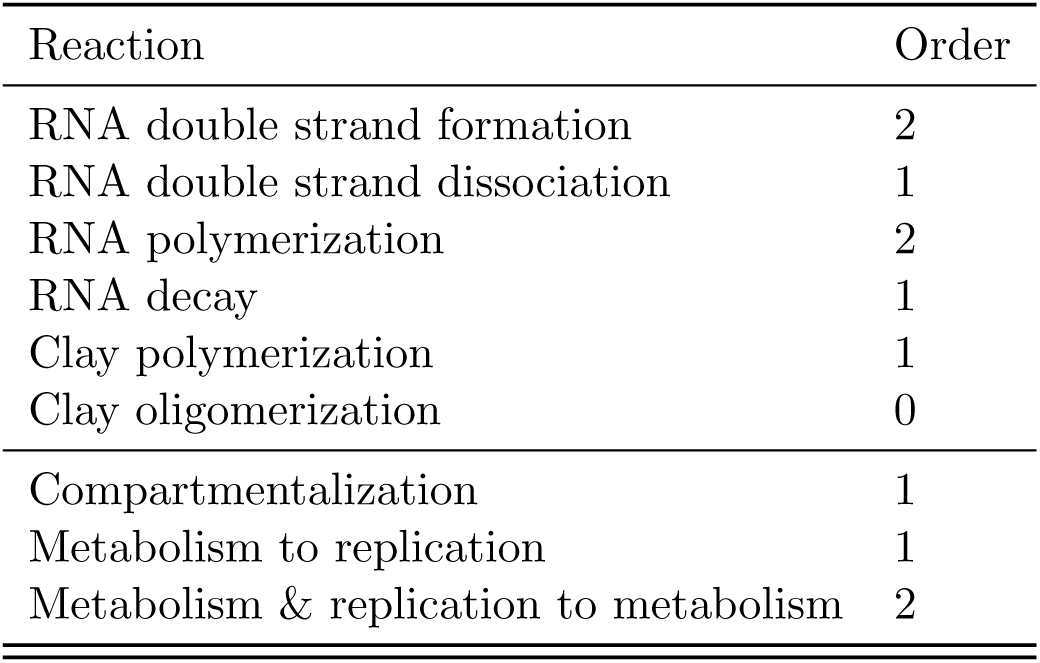
Reactions and orders

| Reaction | Order |
| --- | --- |
| RNA double strand formation | 2 |
| RNA double strand dissociation | 1 |
| RNA polymerization | 2 |
| RNA decay | 1 |
| Clay polymerization | 1 |
| Clay oligomerization | 0 |
| Compartmentalization | 1 |
| Metabolism to replication | 1 |
| Metabolism & replication to metabolism | 2 |

## 3 Results

Consider some initial population of *I* random genomes *X*_0_. The population over time is given by *X*_*t*_ with associated random counting measure *N*_*t*_ on 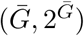. Recall parameters

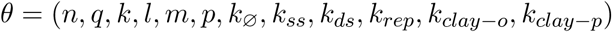

for genome dimension *n*, high-fidelity genome set size *q*, fitness degree *k*, similarity degree *l*, fidelity degree *m*, clay fidelity probability *p*, RNA decay rate *k*_∅_, double-strand dissociation rate *k*_*ss*_, double-strand formation rate *k*_*ds*_, RNA replication rate *k*_*rep*_, and clay oligomerization and polymerization rates *k*_*clay−o*_ and *k*_*clay−p*_. These parameters are summarized in Table 3.

**Table 3:** Model parameters

| $\theta$ | Name | Domain |
| --- | --- | --- |
| $n$ | genome dimension | $\mathbb{N}_{>0}$ |
| $q$ | $R$ probability of subset size 4 for position | $[0, 1]$ |
| $k$ | RNA fitness parameter | $\mathbb{R}_+$ |
| $l$ | RNA similarity parameter | $\mathbb{R}_+$ |
| $m$ | RNA fidelity parameter | $\mathbb{R}_+$ |
| $p$ | clay fidelity probability | $(0, 1]$ |
| $k_{ss}$ | double-strand dissociation rate | $\mathbb{R}_+$ |
| $k_{ds}$ | double-strand formation rate | $\mathbb{R}_+$ |
| $k_{rep}$ | RNA replication rate | $\mathbb{R}_+$ |
| $k_{\emptyset}$ | RNA decay rate | $\mathbb{R}_+$ |
| $k_{clay-o}$ | clay RNA oligomerization rate | $\mathbb{R}_+$ |
| $k_{clay-p}$ | clay RNA polymerization rate | $\mathbb{R}_+$ |

With the parameters governing the reaction rates, different values of these parameters confer different regimes for the system.

### 3.1 Stability: ODEs

We characterize the zeros of the vector field *f* from ODE system (30) and use the eigenvalues of the Jacobian (31) to determine their stability.

#### Theorem 2.

*The ODE system* (30) *for R* = {*x*}, *x* ∈ *E, has a single unstable fixed-point at* [*x*] = 1 *and* [*y*] = 0 *for y* ∈ *G \ x*.

*Proof*. Solving *f* = 0 gives a single solution [*x*] = 1 and [*y*] = 0 for *y* ∈ *G \ x*. For this solution, the eigenvalues of the Jacobian contain no zero values and positive values. Therefore the solution is unstable.

□

It follows from Theorem 2 that for all other initial conditions, the system has no equilibria.

#### Corollary 2.

*For all initial conditions X*_0_ *such that I* = |*X*_0_| *>* 1, *the system is unbounded*.

This confirms the obvious: the system, a replicating network with no death, is almost always an increasing system.

### 3.2 Simulation reaction state

We are interested in the behavior of *v*_*t*_ (25) and *µ*_*t*_ = *v*_*t*_*Q* (28) for RNA polymerization. These reveal the instantaneous information of the system. The structure of *µ*_*t*_ reveals the state of polymerization and is a leading indicator of the population concentrations over time.

#### 3.2.1 Core model with “tent” functions, probable hitting ℙ(*τ*_*v*_(*θ*) *< ∞*) *∼* 1

Take genome dimension *n* = 3 and fitness and similarity parameters *k* = *l* = *−* log(0.01)*/n* and fidelity parameter *m* = *−* log(0.25)*/n*. Set rates *k*_*ss*_ = *k*_*ds*_ = 1 and *k*_*rep*_ = 10 and use the “tent” function for fitness, similarity, and fidelity. Take random *X*_0_ with *I* = |*X*_0_| = 10 and random singleton *R* = {{*x*}} (*q* = 0). We simulate 5 000 reactions, simulation censored at *τ*_*v*_ for *v* = 0.25. Take partition (*H*_*i*_) (29) of *E*. In Figure 1 we plot measures of a typical realization of *X*_*t*_ on (*H*_*i*_) of concentration (Figure 1a), growth (Figure 1b), and *µ*_*t*_ (Figure 1c). Some quantities are plotted on log-log scale, whereas others are plotted on a linear-log scale. These results show that the concentrations are relatively stable for most time, until the high-fidelity manifold is hit. Then the concentration of high-fidelity replicators rapidly increases to exceed 25%. Similarly, Figure 1b shows the growth curves on a log-log scale, where the high-fidelity manifold rapidly increases near the end of the simulation. Figure 1c shows the structure of *µ*_*t*_. Low probability is assigned to polymerization of high-fidelity replicators for most of the reaction time, followed by a large increase near the end of the simulation, where high-fidelity replicators dominate with 56% probability. *Therefore µ*_*t*_ *is a leading indicator of the concentration curve, i*.*e. at simulation end-time, concentration of high-fidelity replicators is 25% and polymerization output is 56%*.

**Figure 1:**
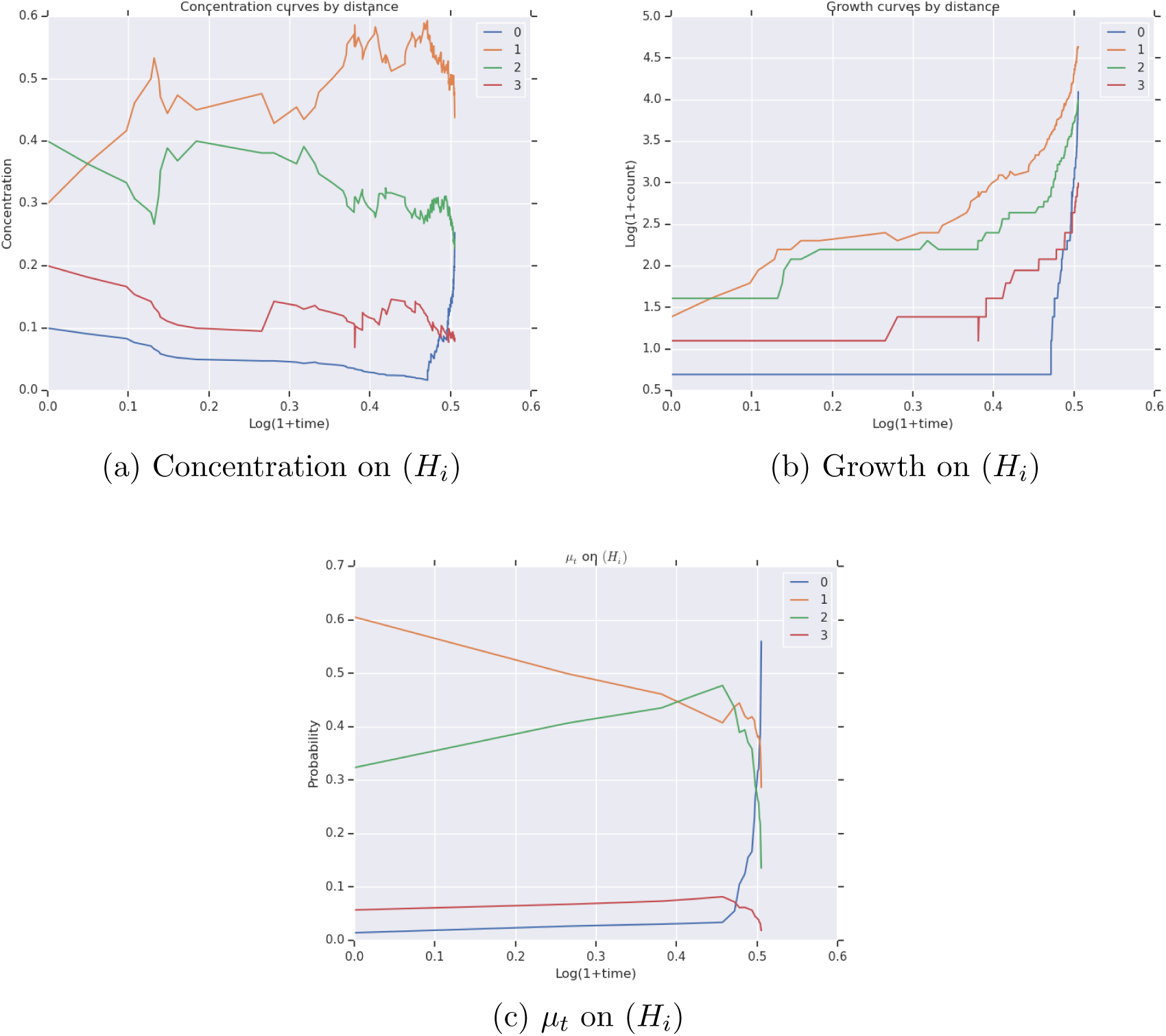
Measures of *X*_*t*_ until *τ*_*v*_ for *v* = 0.25 with *n* = 3 and *k* = *−* log(0.01)*/n* and *I* = |*X*_0_| = 10 and *R* = {{*x*}} and “tent” fitness and similarity functions

#### 3.2.2 Core model with “tent” functions, improbable hitting ℙ(*τ*_*v*_(*θ*) *< ∞*) *∼* 0

We use the same configuration as Section 3.2.1 except for setting *k* = *l* = *−* log(0.1)*/n*. In Figure 2 we plot measures of a typical realization of *X*_*t*_ on (*H*_*i*_) of concentration (Figure 2a), growth (Figure 2b), and *µ*_*t*_ (Figure 2c). The behavior has completely changed: the high-fidelity group ends the simulation with around 6% concentration, only steadily increasing, and never hits. The polymerase output *µ*_*t*_ shows 6%. This indicates that the concentration of high-fidelity replicators is unlikely to increase further, as the population is generally in equilibrium with the polymerase output.

**Figure 2:**
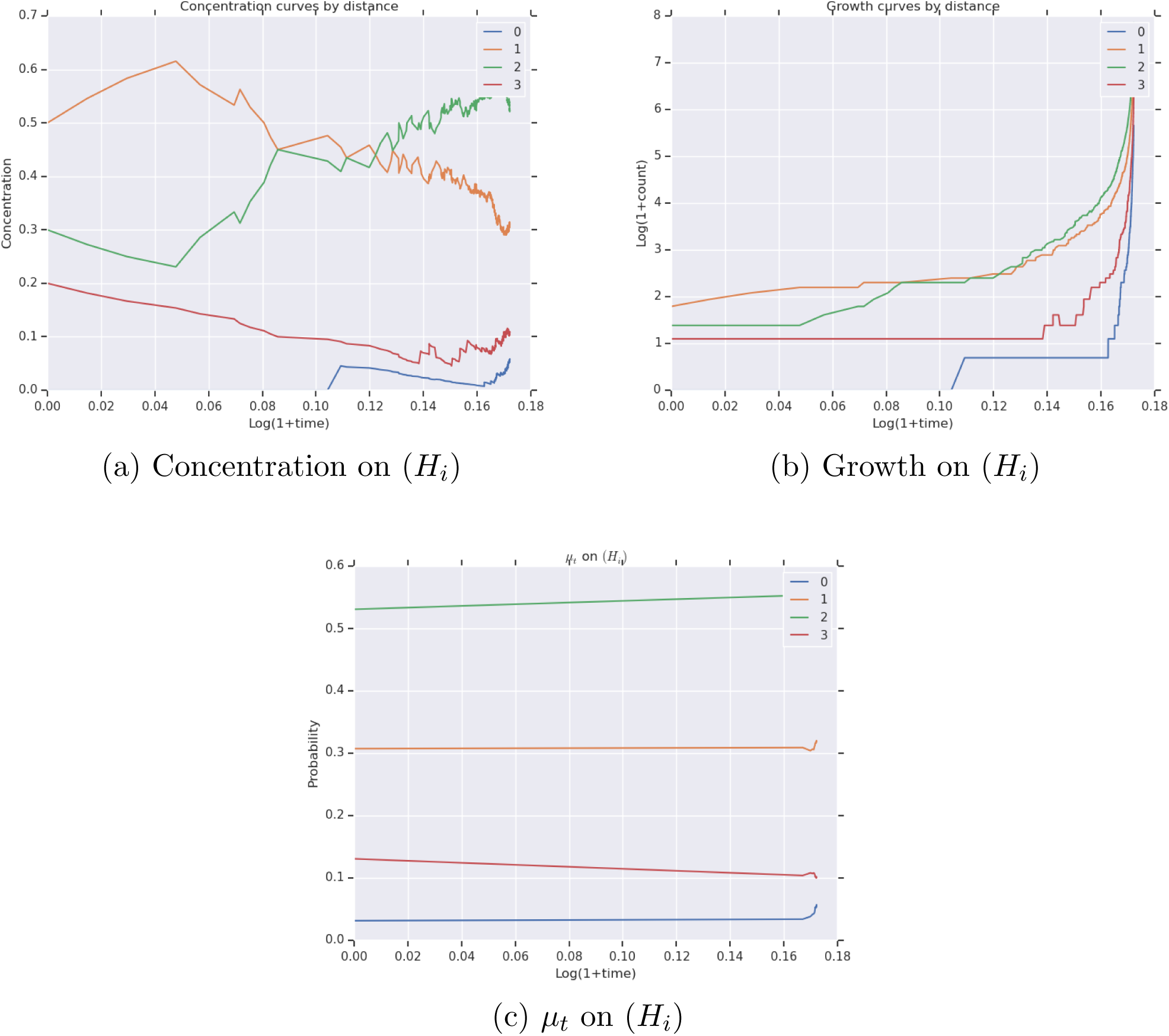
Measures of *X*_*t*_ until *τ*_*v*_ for *v* = 0.25 with *n* = 3 and *k* = *−* log(0.1)*/n* and *I* = |*X*_0_| = 10 and *R* = {{*x*}} and “tent” fitness and similarity functions

#### 3.2.3 Core model with linear functions, improbable hitting ℙ(*τ*_*v*_(*θ*) *< ∞*) *∼* 0

The same configurations for Section 3.2.1 are used, except the fitness, similarity, and fidelity functions are linear. Similar to the “tent” functions, we specify the terminus *k*. Then

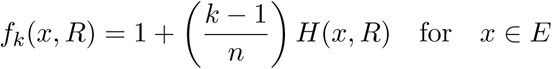

and

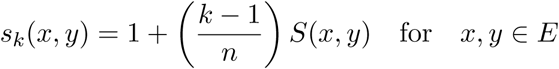

We put *k* = 0.01 for fitness and similarity and *k* = 0.25 for fidelity. We simulate *X*_*t*_ for 5 000 reactions. We find that ℙ(*τ*_0.25_(*θ*) *< ∞*) *∼* 0. In Figure 3 we plot measures of a typical realization of *X*_*t*_ on (*H*_*i*_) of concentration (Figure 3a), growth (Figure 3b), and *µ*_*t*_ (Figure 3c). The simulation ends with high-fidelity concentration of *∼* 5% and polymerase output of *∼* 4%. Therefore the concentration of high-fidelity replicators will continue to decrease. *Linear surfaces are not sufficient to achieve hitting times τ*_*v*_(*θ*) *< ∞ for v* = 0.25, *in contrast to the non-linear “tent” functions*.

**Figure 3:**
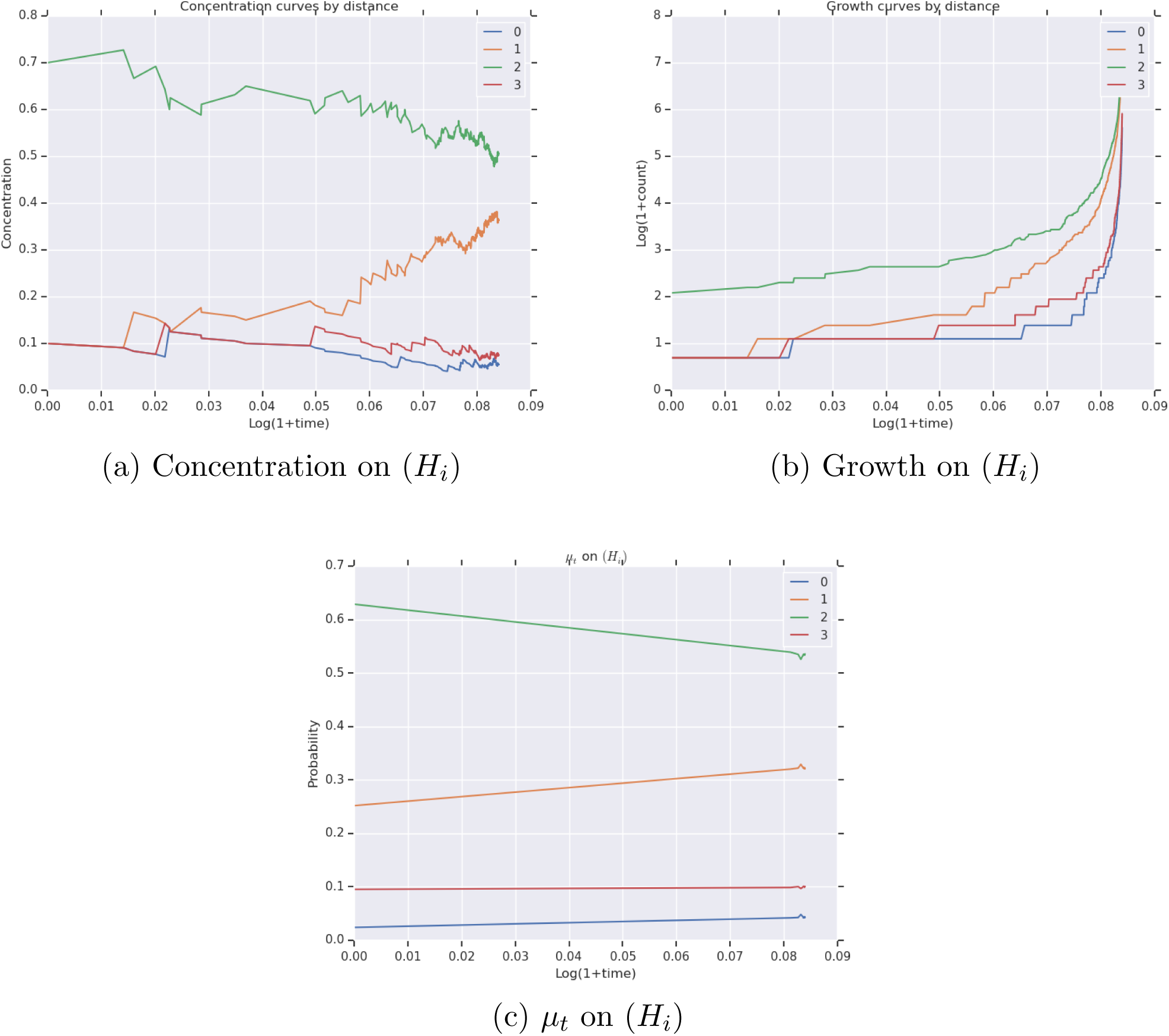
Measures of *X*_*t*_ until *τ*_*v*_ for *v* = 0.25 with *n* = 3 and *k* = *−* log(0.01)*/n* and *I* = |*X*_0_| = 10 and *R* = {{*x*}} and linear fitness and similarity functions

#### 3.2.4 Expanded model with “tent” functions, probable hitting ℙ(*τ*_*v*_(*θ*) *< ∞*) *∼* 1

We consider similar model to the previous subsections and expand it with clay oligomerization rate (of RNA) *k*_*clay−o*_, clay polymerization rate (of RNA) *k*_*clay−p*_, and clay polymerization fidelity *p*. Therefore the full set of variables is given by *θ* = (*n, k*_*ss*_, *k*_*ds*_, *k, k*_*clay−o*_, *k*_*clay−p*_, *p*). The value of *k* and *k*_*clay−p*_ are set such that the replicative mass of each is initialized to 10. This means that RNA and clay polymerization have the same reaction mass at the beginning of the simulation. We set *n* = 3, *k* = *−* log(0.01)*/n, p* = 0.9, and *k*_*ss*_ = *k*_*ds*_ = 1. This is a high hitting regime, i.e. ℙ(*τ*_*v*_(*θ*) *< ∞*) *∼* 1. In Figure 4 we plot measures of a typical realization of *X*_*t*_ on (*H*_*i*_) and additionally the probability of reactions over time. High-fidelity replicators ended the simulation with 25% concentration (Figure 4a) and RNA polymerase output *∼* 62% (Figure 4c), indicating the concentration of high-fidelity replicators will continue to increase. All species exhibit superexponential growth (Figure 4b). Clay polymerization decreases in contribution over time, whereas RNA polymerization increases substantially over time, and RNA double-strand reactions are small and stable (Figure 4d).

**Figure 4:**
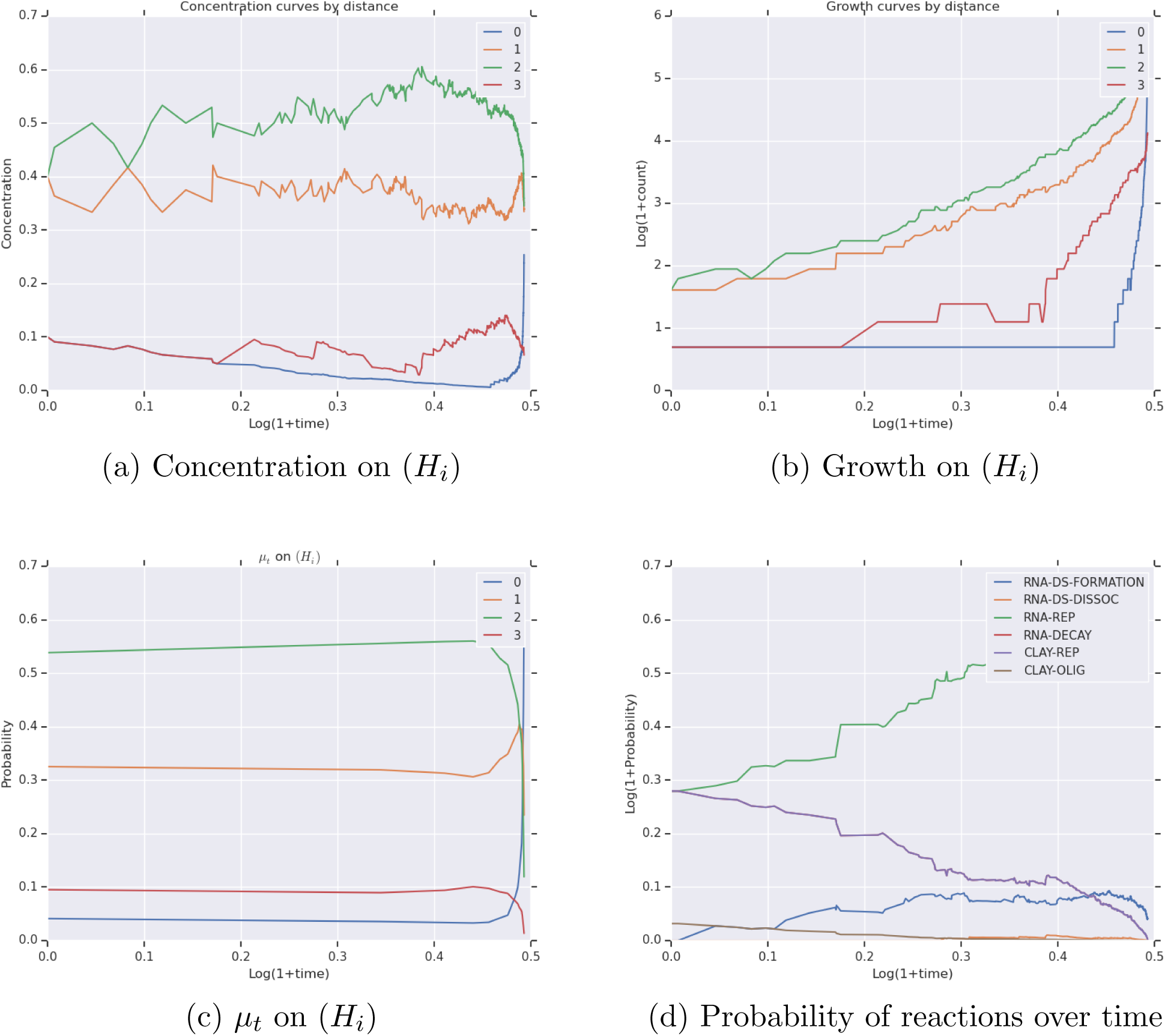
Measures of *X*_*t*_ until *τ*_*v*_(*θ*) for *v* = 0.25 with *n* = 3 and *k* = *−* log(0.01)*/n* and *I* = |*X*_0_| = 10 and *R* = {{*x*}} and *k*_*ss*_ = *k*_*ds*_ = 1 and *p* = 0.9 and *k*_*rep*_, *k*_*clay−p*_ chosen such that the replicative mass of each is 10

### 3.3 Hitting times: functional and survival analysis

We study various models in order of increasing complexity. We examine the hitting time surface *τ*_*v*_(*θ*) in *θ* ∈ Θ, including probability of hitting ℙ(*τ*_*v*_(*θ*) *< ∞*). We begin with the core model with no decay or clay.

#### 3.3.1 Core model, *τ*_*v*_(*θ*) for *v* = 0.1 with *θ* = (*n, k*) and “tent” functions

We are interested in the structure of the hitting time *τ*_*v*_(*θ*) of (19) as a function of *θ*. We use the Weibull-Cox proportional hazard’s model of Table 1 for the hitting time *τ*_*v*_ for *v* = 0.1. Let *θ* = (*n, k*) with genome dimension *n* ∈ {3, 4} and fitness and similarity parameters *k* = *l* = *−* log(*i*)*/n* for *i* ∈ {0.1, 0.05, 0.01, 0.005, 0.001}. Set *m* = *−* log(0.25)*/n* for fidelity probability parameter. For each value of *n*, take random *X*_0_ with *I* = |*X*_0_| = 10 and random singleton *R* = {{*x*}} and fix these for the *k*. We fix the parameters *k*_*ss*_ = *k*_*ds*_ = 1 and set *k*_*rep*_ such that 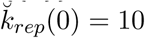 and use the “tent” function for fitness, similarity, and fidelity. We take 10 realizations of *τ*_*v*_(*θ*) for each *θ* ∈ Θ and allocate 5 000 reactions. This gives 100 independent hitting times and up to 500 000 reactions. *The times are comparable because the system is initialized to the same replication mass*.

For the simulations, 66 hitting times are finite. The coefficients positively contribute to hitting, where *γ*_*n*_ ≈ 0.97 and *γ*_*k*_ ≈ 13.29, both with *p*-values less than 0.005. Therefore hittings are strongly positively influenced by the parameters of the fitness and similarity functions and less so by the dimension. Plots of the coefficients and survival and cumulative hazard curves are given in Figure 5. *High survival is found for k large and low survival for k small. Cumulative hazard is highest for k small*.

**Figure 5:**
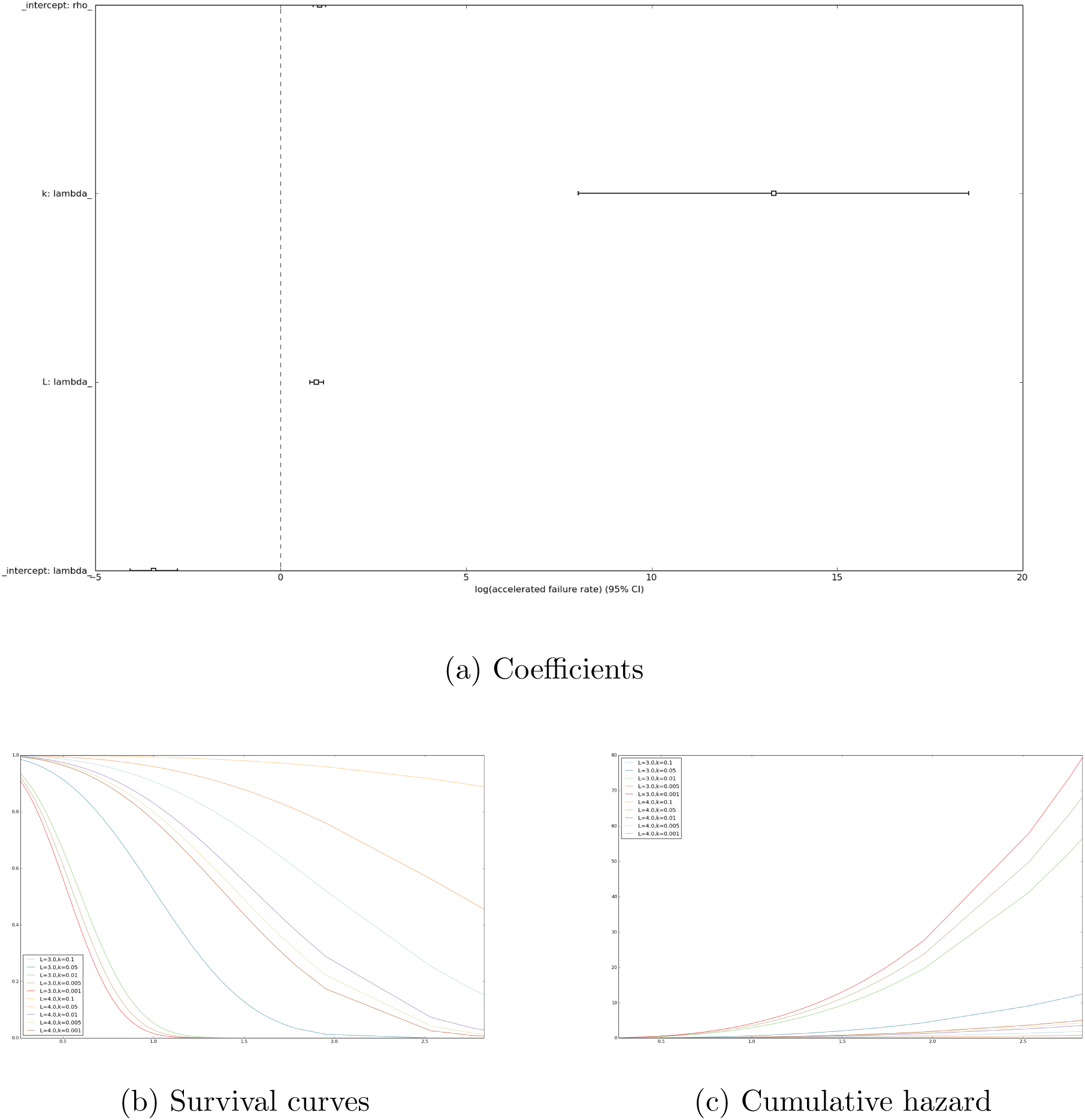
Survival analysis of hitting time *τ*_*v*_ for *v* = 0.1 for core model

We estimate HDMR of the classifier (whether or not hitting time is finite) using all 100 samples. The results are shown in Figure 6. The HDMR explains roughly 80% of variance. 𝕊_*k*_ ≈ 0.69 and 𝕊_*n*_ ≈ 0.06, so *hitting probability is strongly influenced by k and less so by n*. The component functions *f*_*k*_ and *f*_*n*_ are strictly decreasing, where larger *k* results in decreasing hitting probability. These results are consistent with the survival analysis.

**Figure 6:**
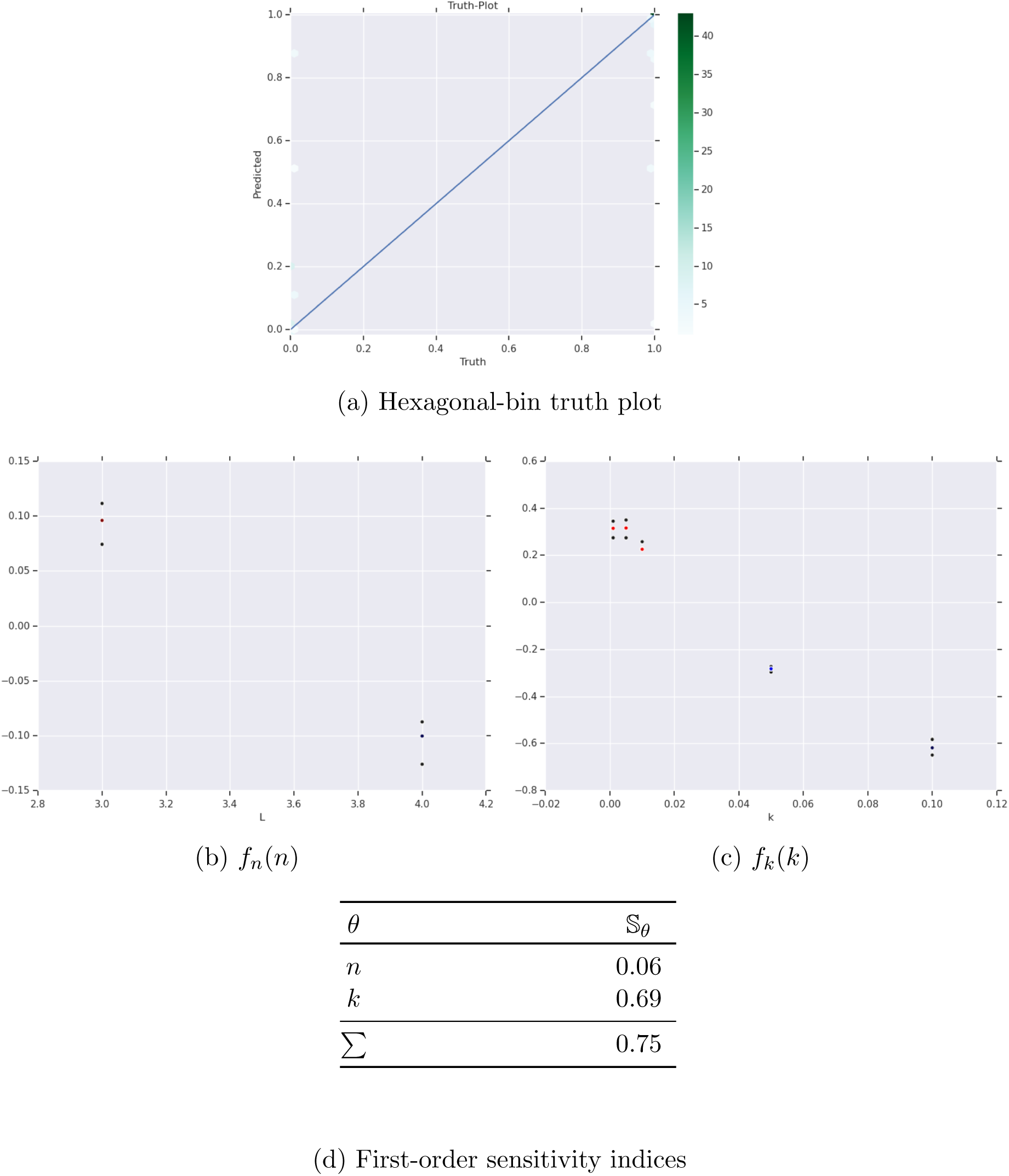
First-order HDMR analysis of ℙ(*τ*_*v*_(*θ*) *< ∞*) for core model

We estimate HDMR of the regressor (hitting time) using the 66 simulations with finite hitting time. The results are shown in Figure 7. The HDMR explains roughly 60% of variance. 𝕊_*n*_ ≈ 0.57 and 𝕊_*k*_ ≈ 0.04, so *genome dimension dominates the hitting time*. Both *f*_*n*_ and *f*_*k*_ are increasing. The HDMR results reveal that conditioning on hitting reverses the roles of *n* and *k*.

**Figure 7:**
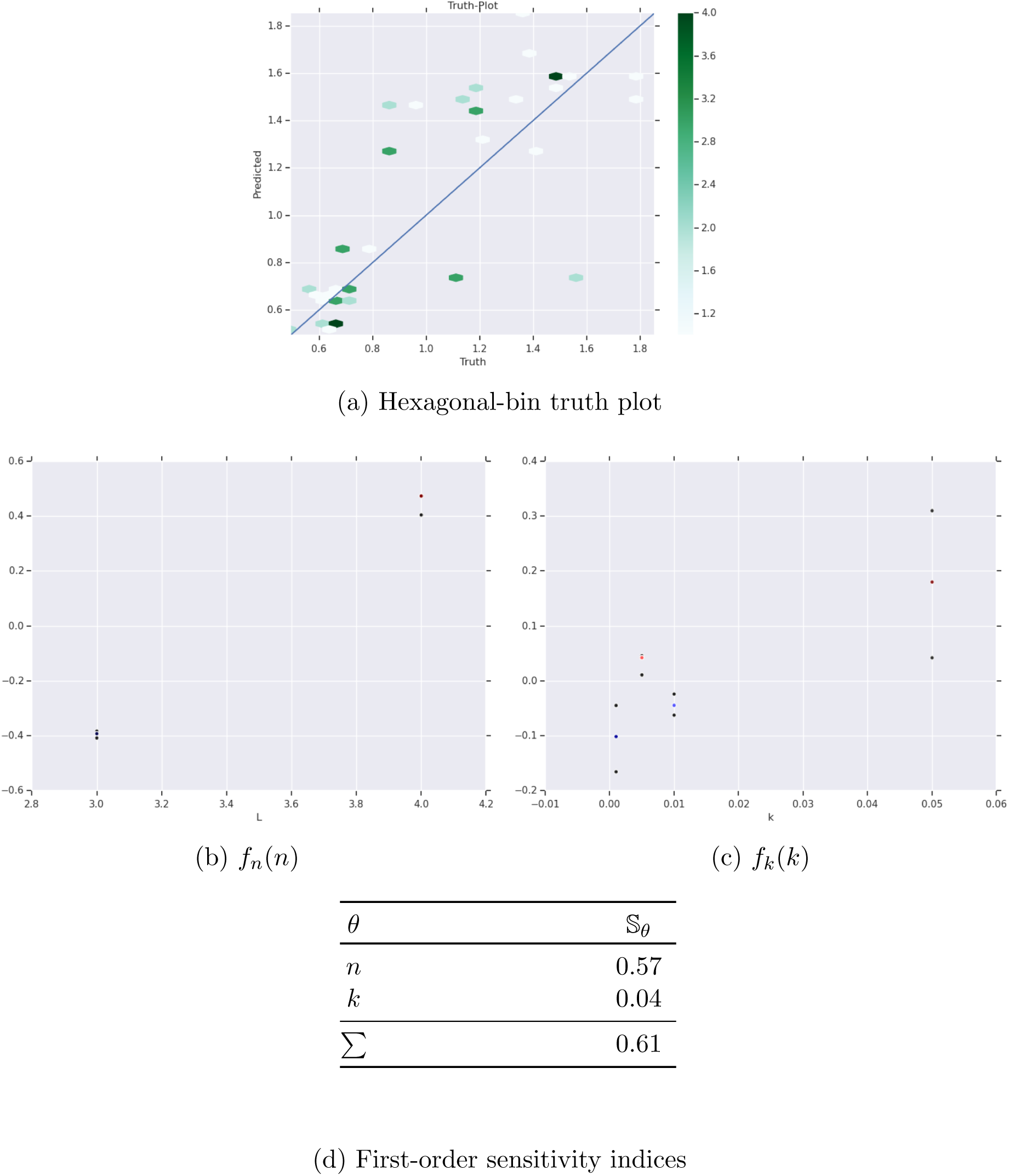
First-order HDMR analysis of *τ*_*v*_(*θ*) *< ∞* for core model

#### 3.3.2 Clay and decay model, *τ*_*v*_(*θ*) for *v* = 0.1 with *θ* = (*n, k, k*_∅_, *k*_*clay−p*_, *p*) and “tent” functions

We expand the model to include clay and decay. We take

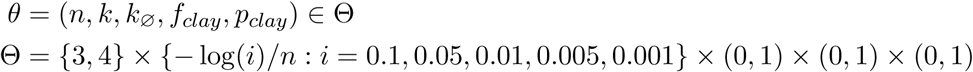

with *k*_*ss*_ = *k*_*ds*_ = *k*_*clay−o*_ = 1. For each *θ* ∈ Θ, (i) we choose random *R* = {*x*}, *x* ∈ *E* and choose random *X*_0_ such that *I* = |*X*_0_| = 10 and *X*_0_ *∩ R* = ∅, that is, the initial population does not reside on the high-fidelity manifold; (ii) we initialize the replicative mass of the system such that 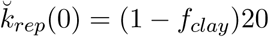 and 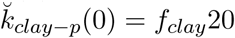; (iii) we sample *τ*_*v*_(*θ*) for *v* = 0.10 *M* = 10 times, each censored by 5 000 reactions, giving *𝒯* (*θ*) of (20). We attain 𝔇 = {(*θ*_*i*_, *𝒯* (*θ*_*i*_)) : *i* = 1,…, 240}. This gives a total of 2 400 simulations.

For the simulations, 1 546 hitting times are finite. The results of fitting the Weibull-Cox model are shown below in Table 4 and Figure 8. The curvature parameter *k* again significantly dominates with a large positive value. All the remaining parameters have values less than one. *n* again is a relatively small positive contributor. The clay fraction *f*_*clay*_ is small and positive and *p* is not significant.

**Table 4:**
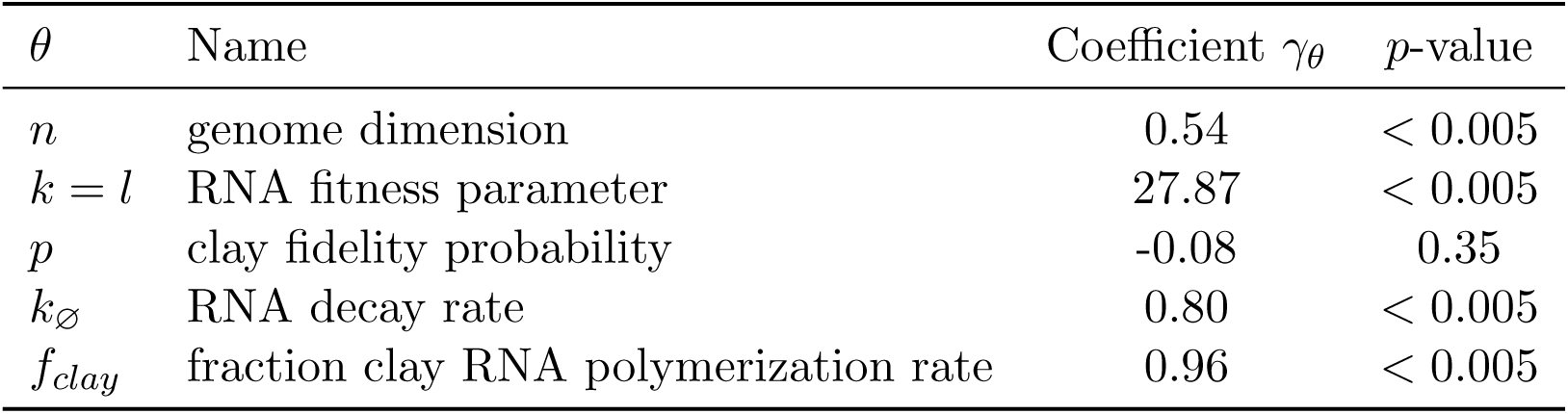
Weibull-Cox model parameters for hitting times of clay and decay model

**Figure 8:**
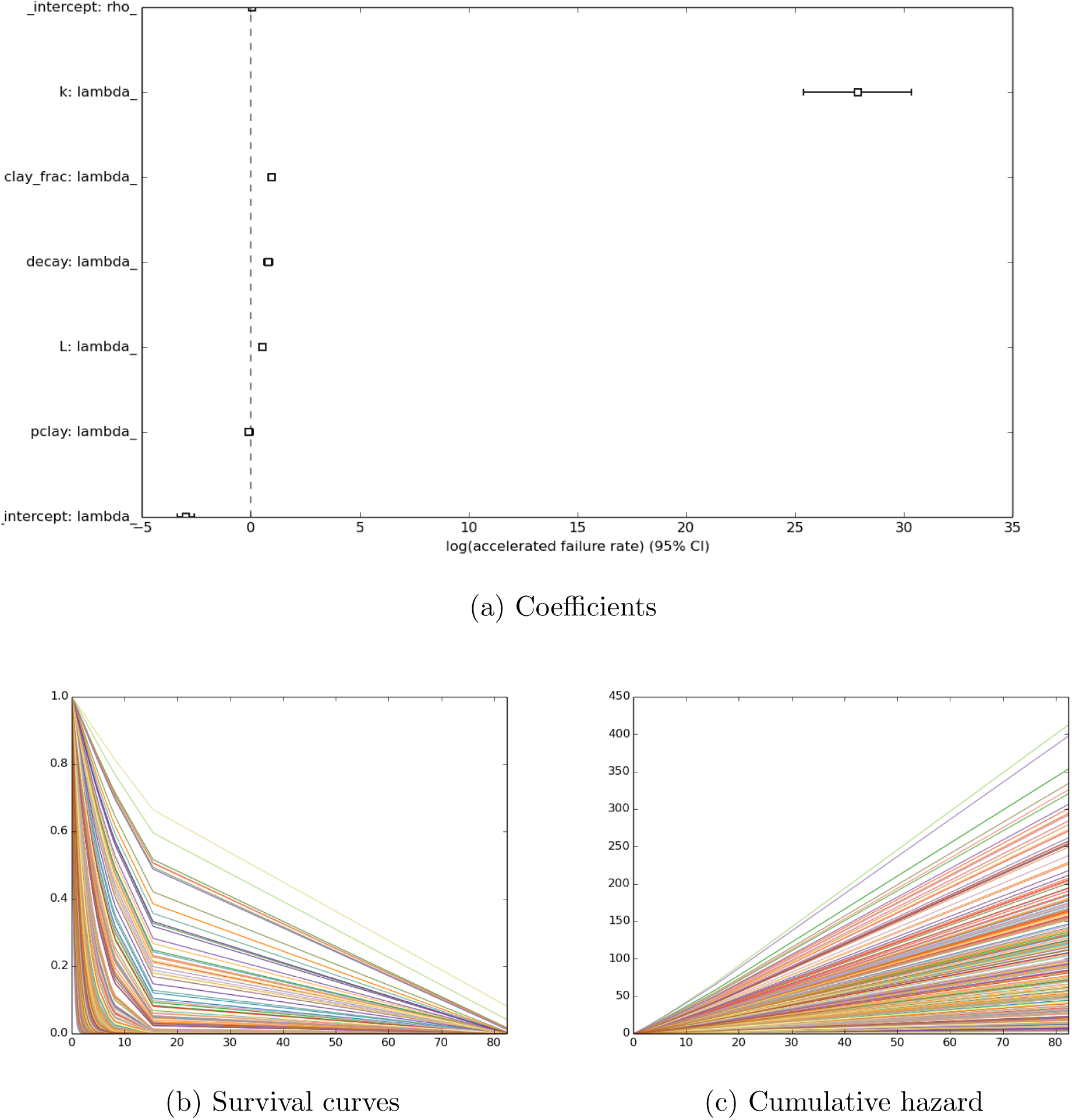
Survival analysis of hitting time *τ*_*v*_ for *v* = 0.1 for expanded model (clay and decay)

We estimate HDMR of the classifier (whether or not hitting time is finite) using all 2 400 samples. Component functions and sensitivity indices are shown below in Figure 9. First-order HDMR captures 74% of explained variance, and second-order captures 4%. *Curvature dominates hitting probability with large sensitivity index* 𝕊_*k*_ ≈ 67%. *f*_*k*_ is a decreasing function, where small values increase and large values decrease hitting probability. Genome dimension *n* has sensitivity index 𝕊_*n*_ ≈ 2%, and *f*_*n*_ is decreasing, where high dimension decreases the probability of hitting. Clay parameter 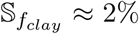, and 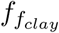 is decreasing, where low-to-medium clay fractions increase and high-clay fractions decrease probability of hitting. The HDMR results are consistent with the Weibull-Cox model.

**Figure 9:**
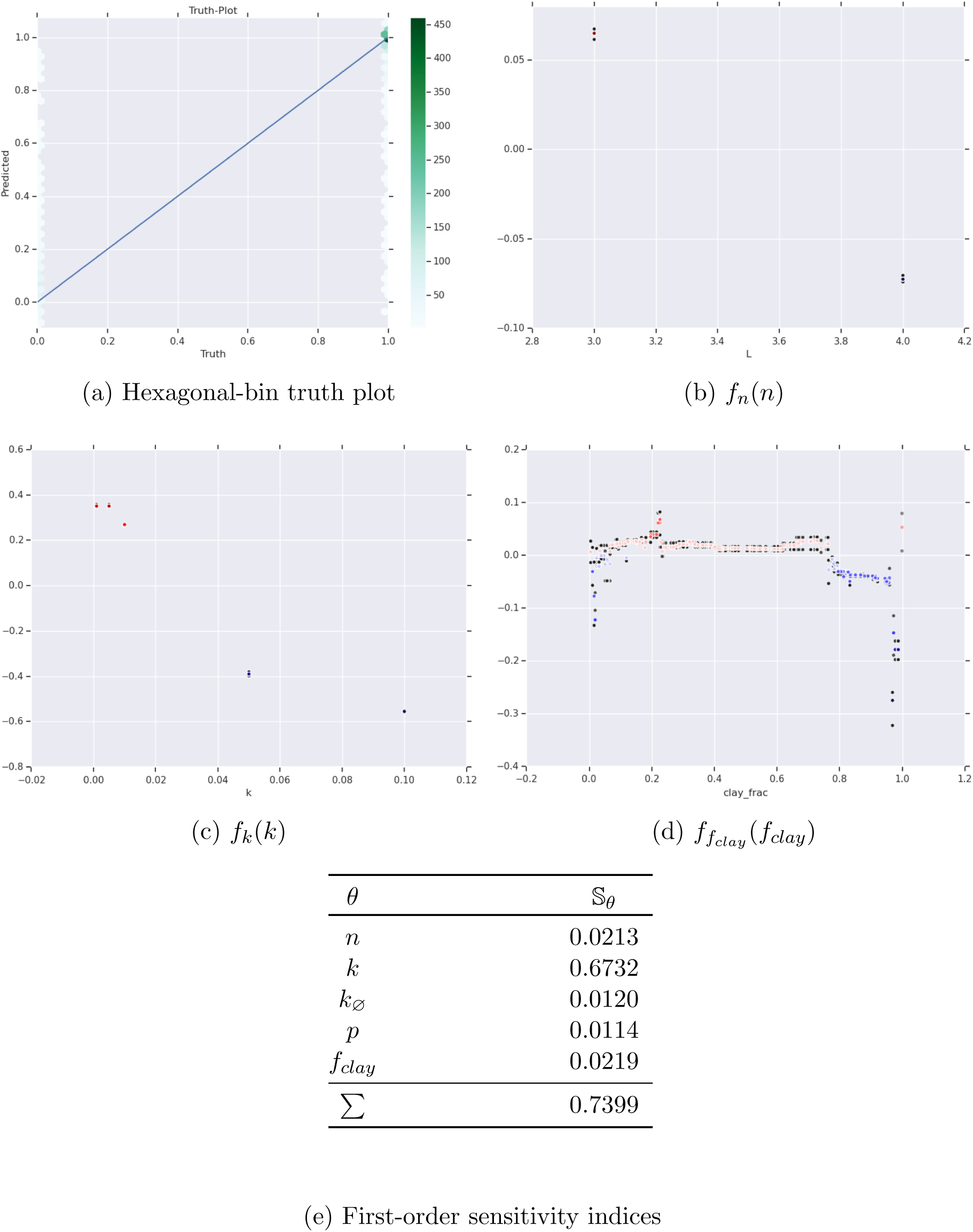
First-order HDMR analysis of ℙ(*τ*_*v*_(*θ*) *< ∞*) for expanded model (clay and decay)

We estimate HDMR of the regressor (hitting time) using the 1 546 simulations with finite hitting time. Component functions and sensitivity indices are shown below in Figure 10. First-order HDMR captures 33% of explained variance, and second-order captures 7%. *In stark contrast to the contributions to the classifier, the parameters k and n are insignificant to hitting time. Instead, the largest sensitivity index is* 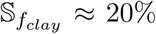. 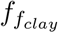 is an increasing function, where small *f*_*clay*_ decreases and large *f*_*clay*_ increases the hitting time. This suggests that high clay-fractions representing first-order reactions increase the hitting time, as clay polymerization has less replicative mass than RNA polymerization, i.e. things go faster with RNA polymerization. The second largest is decay 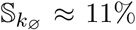. Decay is an increasing function, with sharp increase in hitting times nearby one, i.e. things go slower with large decay resulting in increased hitting time.

**Figure 10:**
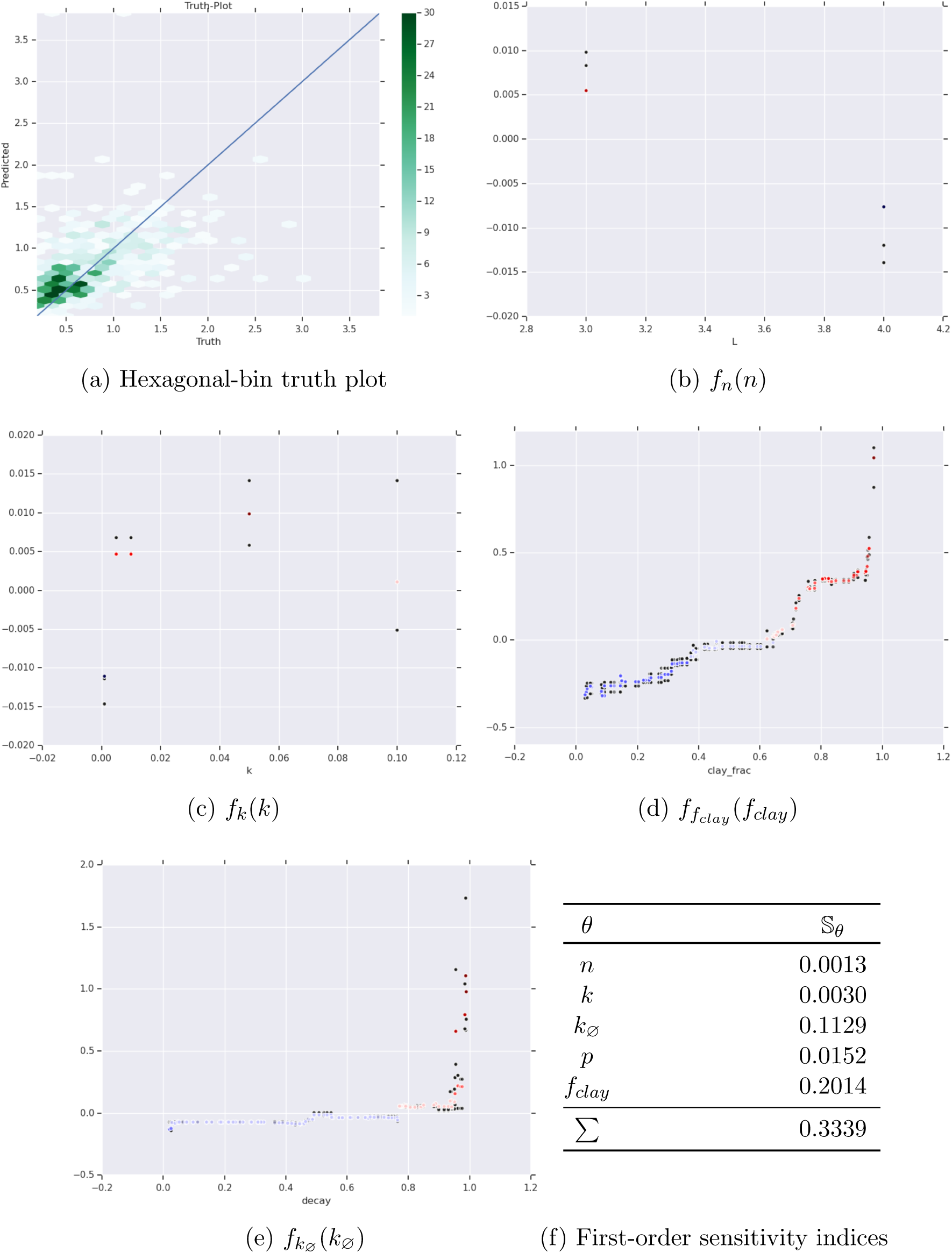
First-order HDMR analysis of *τ*_*v*_(*θ*) *< ∞* for expanded model (clay and decay)

### 3.4 Reaction cardinality

In this section we study, instead of the hitting time *τ*_*v*_(*θ*), the hitting reaction number *ϖ*_*v*_(*θ*) for *v* = 0.1. We study the core model with *θ* = (*n, k*). We uniformly sample *n* ∈ {3, 4, 5} and *k* ∈ {0.0001, 0.0005, 0.001, 0.005, 0.01, 0.05, 0.1} and form data 𝔇 = {(*θ*_*i*_, *rv*(*θ*_*i*_)) : *i* = 1,…, 1 200}. We set *k*_*ss*_ = *k*_*ds*_ = 1 and initialize *k*_*rep*_ such that the initial replicative mass is 10. For each *θ* ∈ Θ, we randomly choose *X*_0_ with *I* = |*X*_0_| = 10 and singleton *R* such that *X*_0_ *∩ R* = ∅.

𝔇 contains 781 hitting events. We denote these 𝔇^*\**^. We form a first-order HDMR on *rv*(*θ*) using 𝔇^*\**^. The HDMR truth-plot, component functions, and sensitivity indices are shown below in Figure 11. First-order HDMR captures approximately 31% of variance. Both component functions *f*_*n*_ and *f*_*k*_ have similar sizes, with *f*_*n*_ somewhat larger than *f*_*k*_. *f*_*n*_ is essentially an increasing linear function of *n*, and *f*_*k*_ is a generally an increasing function. *These results suggest that genome dimension and curvature influence the hitting reaction*. Larger genomes and flatter curvature increase the hitting reaction.

**Figure 11:**
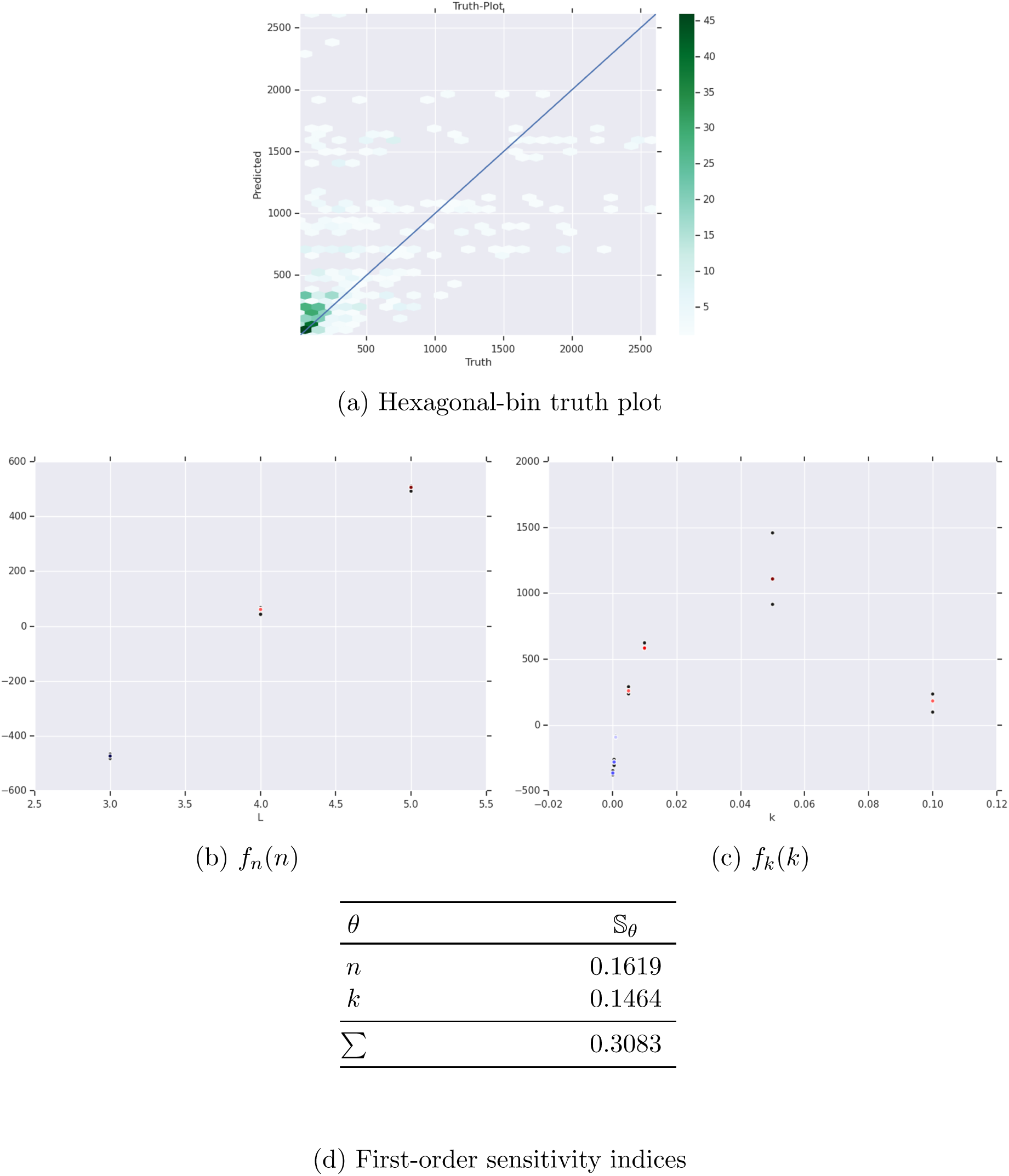
First-order HDMR analysis of *ϖ*_*v*_(*θ*) < ∞ for core model and *v* = 0.1

### 3.5 Compartmentalization

Compartmentalization has a direct effect on the calculation of the reaction rates, specifically replication, by computing only a subset of the reactions in 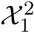. Put

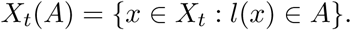

For micelle region *A* ∈ 𝔐 we have that

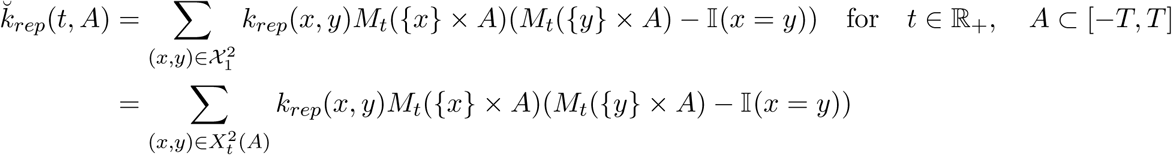

and total replicative mass

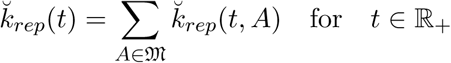

As 𝔐 increases in size over time, the number of partitions grows, and

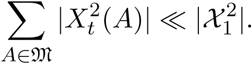

Therefore the replicative mass will be reduced with M, and the system evolves less quickly. *This suggests that compartmentalization should occur after replication. If it co-occurs with the RNA search, then it slows the system down*.

## 4 Conclusions

Origins of life is a fascinating problem. The wonderful complexity of extant life follows from origins. The distribution of life in the universe is tied to origins.

In this article, we have attempted to “peek” into the problem by concentrating on the RNA world hypothesis, studying hitting times of high-fidelity replicators. We develop fitness, similarity, and fidelity functions as landscapes for a mathematical model of replicating RNA molecules at the genome level and observe hitting times through simulation studies. We draw attention to the distinction between the probability of hitting ℙ(*τ* (*θ*) *< ∞*) and the hitting time *τ* (*θ*) *< ∞*.

In terms of mathematical set-up, we interpret the reactions as measure-kernel-functions. Each reaction is identified to and fully encoded by a probability transition kernel. This applies as well to other ‘privileged’ functions, with corresponding transition kernels, although these in particular remain abstract and are generally outside the intended scope of this article. The reactions take place in some domain, whereby all molecules may interact. We note that, in reality, molecules have limited diffusion, and this effectively breaks the reaction domain into independent subdomains above some length scale, i.e. molecules are more likely to react with their neighbors. Therefore we assume our reaction volume is sufficiently small such that all molecules may participate in the reactions.

We find that *non-linear landscapes* are *necessary* for hitting: linear landscapes are insufficient. For non-linear landscapes, we find that the *probability of hitting* is dominated by *curvature* and that *hitting times* are dominated by *genome dimension*. These results suggest that the landscapes in nature are non-linear with high curvature, and that the hitting time for high-fidelity replicators is an increasing function of genome dimension. When clay and decay are added to the model, hitting probability is again dominated by curvature, and clay and decay are relatively insignificant. However clay and decay significantly contribute to the hitting times, in both cases increasing hitting times.

In terms of the relation between replication and compartmentalization and metabolism, we find that compartmentalization slows system dynamics and should occur after the RNA search is concluded. Therefore compartmentalization is identified to a single transition kernel *Q*_*c*_, tied to genomic adaptation. It is possible to argue that because the system takes place inside some domain, i.e. a compartment, compartmentalization is a necessary condition. Metabolism is thought to be identified to production of precursors to RNA synthesis, leading to replication identification, followed by genomic adaptation to metabolism. Metabolism is thus defined through two transition kernels, *Q*_*m*_ and *Q*_*e*_. The independence of these transition kernels can be called into question, and it is quite possible that general transition kernels on the full product spaces may be necessary to satisfactorily explain origins. That is, all three ‘privileged’ functions may have co-evolved.

If we let *f* ∈ ℰ_+_ be a fitness function, then the fitness value *J*(*µ*_*t*_) = *µ*_*t*_*f* is the expected value of the fitness function with respect to the probability measure *µ*_*t*_ = *v*_*t*_*Q*. In OptiEvo theory, *J*(*µ*_*t*_) is studied as a function of the population *X*_*t*_ on (*E, ℰ*) (Feng et al., 2012). OptiEvo assumes that the set of all probability measures {*µ*_*t*_} is convex and that *X*_*t*_ has sufficient flexibility such that *J*(*µ*_*t*_) may be explored around *µ*_*t*_. Then OptiEvo predicts that *J*(*µ*_*t*_) has global maxima on (*E, ℰ*) and that these form a connected level-set of genomes with the same fitness value. Both predictions hold for our model. The first prediction is satisfied by zero distance in fitness and similarity functions for high-fidelity genomes. The second prediction is implied by the high-fidelity set being a singleton or a product-space construction. The main point of contention is whether *X*_*t*_ has sufficient flexibility in exploring *J*(*µ*_*t*_) around *µ*_*t*_. This has direct bearing on the structure of *Q*: if *X*_*t*_ is infexible, then *Q* is constrained to certain subspaces of (*E, ℰ*), i.e. not all transitions are possible.

For future work, the model could be extended to the space of genomes of lengths up to *n*, (*E*^*\**^, *ℰ*^*\**^) or even the space of genomes of all lengths, where distance and similarity functions would utilize a more general string distance metric, e.g., Levenshtein distance. We note that the size of (*E*^*\**^, *ℰ*^*\**^) is not much larger than (*E, ℰ*). The aforementioned transition kernels *Q*_*c*_, *Q*_*m*_, and *Q*_*e*_ could be studied. More general models for polymerization *Q* based on sequence-specific schemata could be developed, using the structure of the Poisson-binomial distribution. It would be interesting to study lipid-RNA and metabolism-RNA interactions and equip the system with the ability to append nucleotides to their genomes to form genes, such as storing useful information for the replication channel. For example, genomes could become adapted relative to schemata that code for reproductively useful functions, e.g., a “primordial shield” hypothesis of lipids that interact with specific genomes which confers protection against decay through shielding.

## A Discrete probability space

We define some concepts related to the space (*E, ℰ*). The discrete probability measure *v* on (*E, ℰ*) is defined by

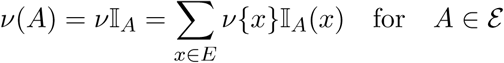

where *v*{*x*} is the probability mass at the point *x* ∈ *E*. For the collection of positive *ℰ* -measurable functions *ℰ*_+_ we have

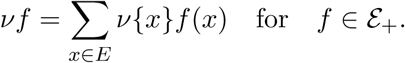

## B Other fitness functions

Another fitness function can be defined using polynomials, such as lines, quadratics, etc, in terms of *k* ∈ ℕ_*>*0_

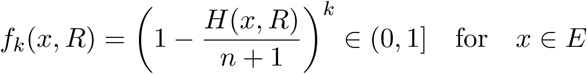

or

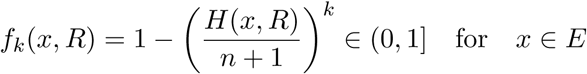

Yet another surface is using a sigmoid function. Put

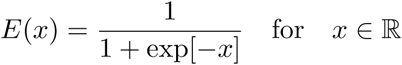

We have fitness for *k* ∈ (0, *∞*)

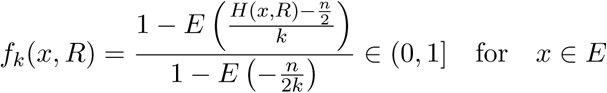

## C Measure-kernel-function

We recall a few facts about transition kernel *Q. Q* defines a function

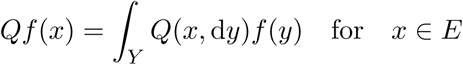

that is in *𝒳*_+_ for every function *f* ∈ *𝒴*_+_. For every probability measure *v*_*t*_ on (*E, ℰ*) the quantity *µ*_*t*_ = *v*_*t*_*Q* defines a probability measure on (*Y, 𝒴*) as

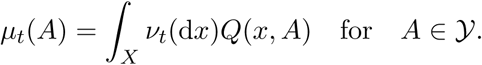

For every probability measure *v*_*t*_ on (*X, 𝒳*) and function *f* ∈ *𝒴*_+_ we have that

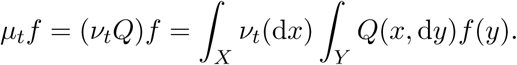

Here the spaces are discrete, i.e.

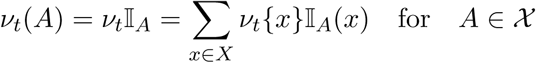

where *v*_*t*_{*x*} is the probability mass at the point *x* ∈ *X*.

## D Approximate reaction rates

One approach to reducing the computational complexity of 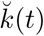 is to approximate the sums using Monte Carlo. Define random variables 𝔛_1_ *∼* Uniform(*𝒳*_1_) and 𝔛_2_ *∼* Uniform(*𝒳*_2_). Let {𝔛_1*i*_} and {𝔛_2*i*_} be independencies of such random variables. Given *N* random samples of 𝔛_1_, the first reaction rate becomes

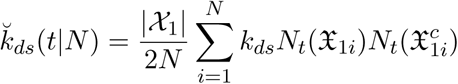

whose expected value is approximated using *M* realizations,

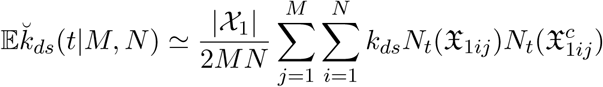

requiring a total of *MN* evaluations. In a similar manner, the second reaction rate is

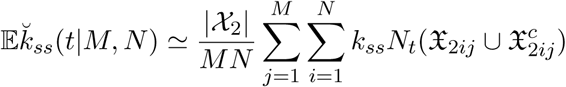

and putting 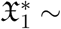 Uniform(𝒳_1_) we have the third reaction

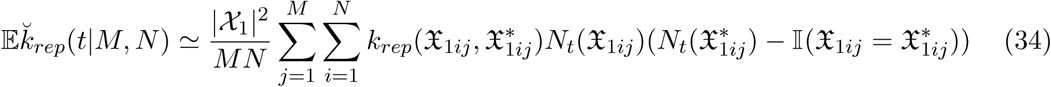

We refer to SSA simulation with Monte Carlo approximate reaction rates as Monte Carlo Approximate SSA, or MCASSA.

### MCASSA performance analysis

At any time point in time, the reaction rates are determined from the population *X*_*t*_. We focus on the replication rate. We study the Monte Caro estimator performance by varying the population *X*_*t*_ of single-stranded genomes of size *m*, genome dimension *n, q* (which is used to construct *R*), *k* (fitness degree), and *l* (similarity degree). We take *θ* = (*n, q, m, k, l*) ∈ Θ = {1,…, 10^2^} × (0, 1) × {1,…, 10^3^} × {1,…, 6} × {1,…, 6} with uniform *v* = *v*_*n*_*v*_*q*_*v*_*m*_*v*_*k*_*v*_*l*_. We generate input data {*θ*_*i*_}. For each *θ*_*i*_ ∈ Θ, we attain *R*_*i*_. Then, given *θ*_*i*_ and *R*_*i*_, we estimate (34) 100 times using Monte Carlo with *MN* = 10^3^ and compute the mean to standard deviation (signal-to-noise ratio) 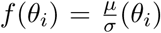 of the samples. *f* encodes the performance or efficiency of the reaction rate estimator. This procedure is repeated 10^3^ times to generate input-output data *𝒟* = {(*θ*_*i*_, *f* (*θ*_*i*_)) : *i* = 1,…, 10^3^}. From *𝒟* we estimate first-order HDMR of *f*

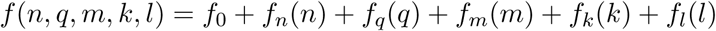

using degree 5 polynomials in each variable and then compute the sensitivity indices 𝕊_*n*_, 𝕊_*q*_, 𝕊_*m*_, 𝕊_*k*_, and 𝕊_*l*_. Truth plots and first-order component functions and sensitivity indices are shown below in Figure 12. First-order HDMR captures *∼* 74% of variance. The largest contributor is similarity degree *l* with *∼* 48% of variance, where increasing *l* decreases the signal-to-noise ratio. The next largest contributor is genome dimension *n* with *∼* 13% of variance, where increasing *n* increases the signal-to-noise ratio. The next largest contributor is the fraction *q* with *∼* 9% of variance, where increasing *q* increases the signal-to-noise ratio. The next largest contributor is fitness degree *k* with *∼* 4% of variance, where increasing *k* decreases the signal-to-noise ratio. The smallest contributor by far is population size *m* with *∼* 0.04% of explained variance. This shows that the estimator performance is generally independent of the population size.

**Figure 12:**
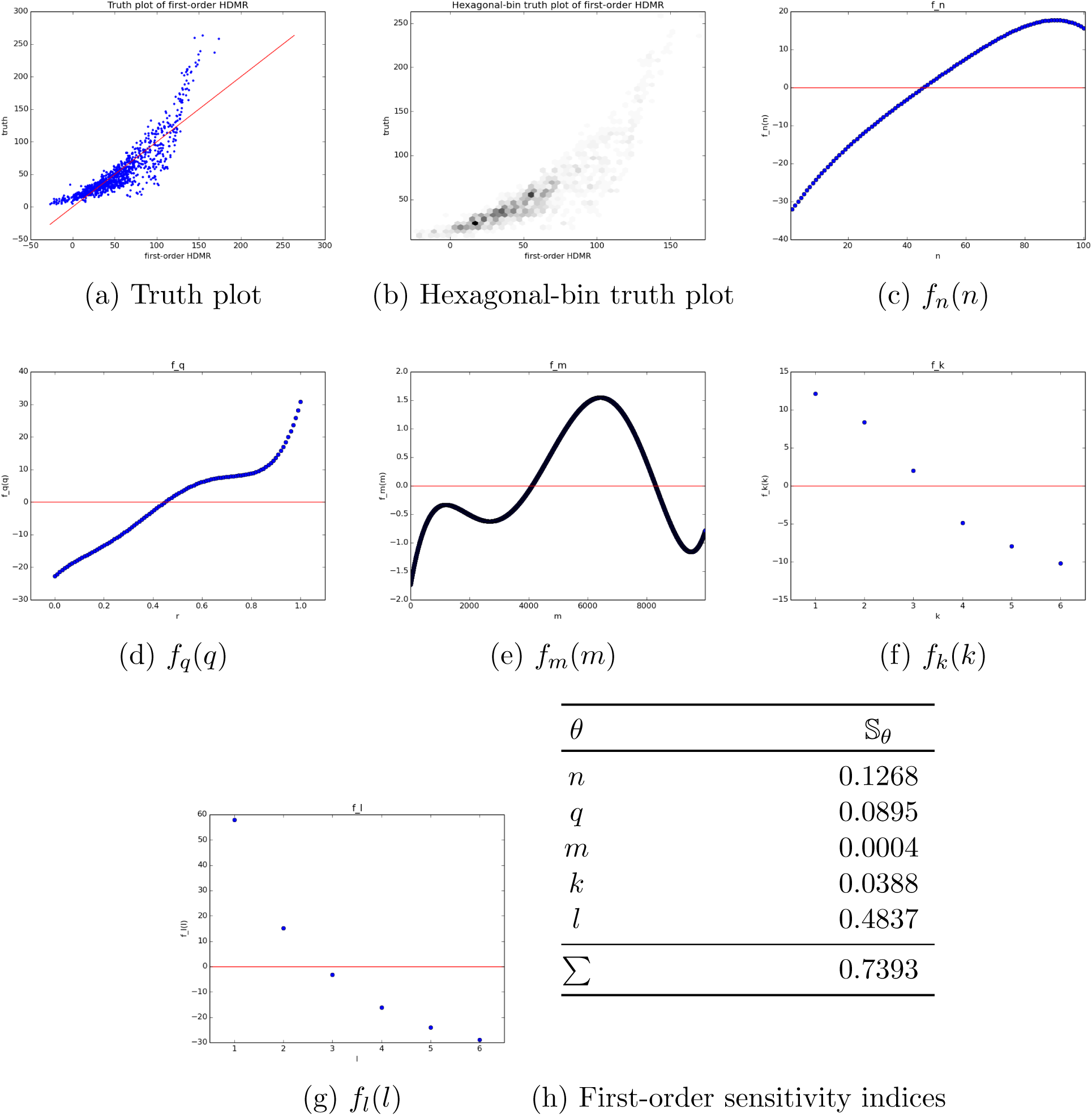
First-order HDMR analysis of *f* (*n, r, m, k, l*)

## E High dimensional model representation

Suppose we have a real-valued square-integrable function *f* (*x*) ∈ *L*^2^(*E, ℰ, v*) with *E* = ℝ^*n*^, 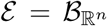, *v* = ∏_*i*_ *v*_*i*_, and *x* = (*x*_1_,…, *x*_*n*_) ∈ *E*. The {*v*_*i*_} may be diffuse (continuous) and/or discrete. Put *B* = {1,…, *n*}. We would like to decompose *f* into orthogonal function subspaces {*f*_*u*_ : *u ⊆ B*} (projections) in such a way that each projection on an input subspace *f*_*u*_ maximizes variance and across subspaces retrieves total variance, i.e. *f* = ∑_*u⊆B*_ *f*_*u*_ and 𝕍*arf* = ∑_*u⊆B*_ 𝕍*arf*_*u*_. The solution to this problem in the retrieval of {*f*_*u*_ : *u ⊆ B*} is known as *high dimensional model representation* (HDMR) or functional ANOVA expansion and for *f* is written as

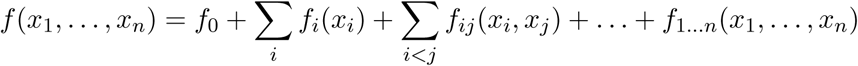

where the {*f*_*u*_ : *u ⊆ B*} are called *component functions*. For independent inputs, the component functions are mutually orthogonal and, aside from the constant component function *f*_0_ = 𝔼*f* (order zero), have zero mean 𝔼*f*_*u*_ = 0 for all non-empty (2^*n*^*−*1) subspaces, where

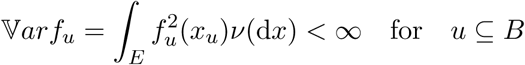

and

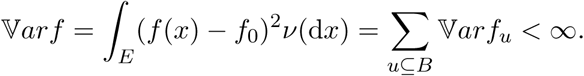

A key principle of HDMR is that the expansion for most *f* may be truncated at low order *T* ≪ *n* in a *T* -order HDMR,

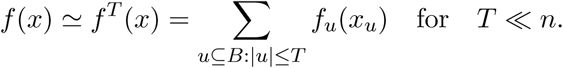

HDMR is often used in *global sensitivity analysis* to assess input-output correlations at various orders, where the variances are normalized to define *sensitivity indices*

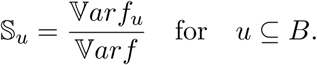

When the inputs are correlated *v ≠∏*_*i*_ *v*_*i*_, then the component functions may still be uniquely recovered under hierarchical orthogonality, the variance decomposes

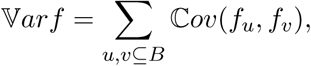

where

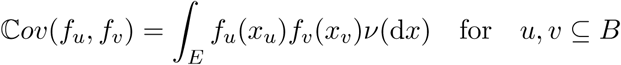

and the sensitivity indices generalize to *structural* and *correlative* sensitivity indices (Li and Rabitz, 2012), defined respectively as

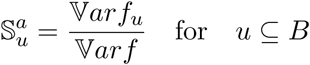

and

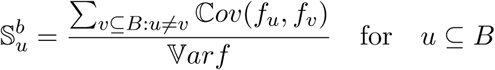

with *total* sensitivity index

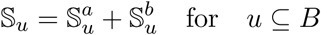

where ∑_*u⊆B*_ 𝕊_*u*_ = 1. We use the total sensitivity index as a measure of variable importance and the component functions as profiles of output dependence on the input subspaces.

## F Reliability definitions

Given a failure distribution *f* and reliability (survival) distribution *R*, we give some relations: the cumulative failure distribution is defined as

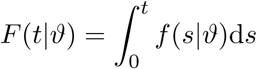

where

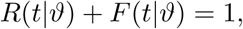

the hazard rate *h*(*t*|*ϑ*) is defined as

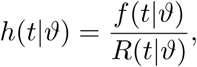

the cumulative hazard is defined as

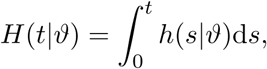

and we have reliability expressed in terms of the cumulative hazard

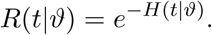

Another useful quantity is the mean residual life

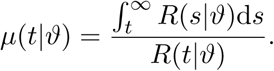

## G Declarations

### Author contributions

All authors contributed to the study conception and design. Material preparation, data generation and analysis were performed by Caleb Bastian. The first draft of the manuscript was written by Caleb Bastian and all authors commented on previous versions of the manuscript. All authors read and approved the final manuscript.

### Ethics approval

Not applicable

### Consent to participate

Not applicable

### Consent for publication

Not applicable

## References

Attwater, J., Wochner, A., and Holliger, P. (2013). In-ice evolution of rna polymerase ribozyme activity. Nature chemistry, 5(12):1011–1018.

Bornholt, J., Lopez, R., Carmean, D. M., Ceze, L., Seelig, G., and Strauss, K. (2016). A dna-based archival storage system. In Proceedings of the Twenty-First International Conference on Architectural Support for Programming Languages and Operating Systems, ASPLOS ‘16, pages 637–649, New York, NY, USA. Association for Computing Machinery.

Cairns-Smith, A. G. (1987). Genetic Takeover and the Mineral Origins of Life. Cambridge University Press.

Cech, T. R. (2000). The ribosome is a ribozyme. Science, 289(5481):878.

Coveney, P. V., Swadling, J. B., Wattis, J. A. D., and Greenwell, H. C. (2012). Theory, modelling and simulation in origins of life studies. Chemical Society Reviews, 41(16):5430–5446.

Davidson-Pilon, C. (2019). lifelines: survival analysis in python. Journal of Open Source Software, 4(40):1317.

Diener, T. O. (1971). Potato spindle tuber “virus”. iv. a replicating, low molecular weight rna. Virology, 45(2):411–428.

Dyson, F. (1999). Origins of Life. Cambridge University Press, Cambridge.

Ehrenfreund, P., Rasmussen, S., Cleaves, J., and Chen, L. (2006). Experimentally tracing the key steps in the origin of life: The aromatic world. Astrobiology, 6(3):490–520.

Eigen, M. and Schuster, P. (1977). A principle of natural self-organization. Naturwissenschaften, 64(11):541–565.

Ertem, G. and Ferris, J. P. (1997). Template-directed synthesis using the heterogeneous templates produced by montmorillonite catalysis. a possible bridge between the prebiotic and rna worlds. Journal of the American Chemical Society, 119(31):7197–7201.

Feng, X., Pechen, A., Jha, A., Wu, R., and Rabitz, H. (2012). Global optimality of fitness landscapes in evolution. Chemical Science, 3(3):900–906.

Ferris, J. P. (2006). Montmorillonite-catalysed formation of rna oligomers: the possible role of catalysis in the origins of life. Philosophical transactions of the Royal Society of London. Series B, Biological sciences, 361(1474):1777–1786.

Gilbert, W. (1986). Origin of life: The rna world. Nature, 319(6055):618–618.

Gillespie, D. T. (1977). Exact stochastic simulation of coupled chemical reactions. The Journal of Physical Chemistry, 81(25):2340–2361.

Hays, L. (2015). Nasa astrobiology strategy. Technical report, National Aeronautics and Space Administration.

Hordijk, W. and Steel, M. (2004). Detecting autocatalytic, self-sustaining sets in chemical reaction systems. Journal of Theoretical Biology, 227(4):451–461.

Joyce, G. F. (2002). The antiquity of rna-based evolution. Nature, 418(6894):214–221.

Kitadai, N. and Maruyama, S. (2018). Origins of building blocks of life: A review. Geoscience Frontiers, 9(4):1117–1153.

Kunin, V. (2000). A system of two polymerases –a model for the origin of life. Origins of life and evolution of the biosphere, 30(5):459–466.

Kvenvolden, K., Lawless, J., Pering, K., Peterson, E., Flores, J., Pon-Namperuma, C., Kaplan, I. R., and Moore, C. (1970). Evidence for extraterrestrial amino-acids and hydrocarbons in the murchison meteorite. Nature, 228(5275):923–926.

Lanier, K. A. and Williams, L. D. (2017). The origin of life: Models and data. Journal of Molecular Evolution, 84(2).

Li, G. and Rabitz, H. (2012). General formulation of hdmr component functions with independent and correlated variables. Journal of Mathematical Chemistry, 50(1):99–130.

Martin, W., Baross, J., Kelley, D., and Russell, M. J. (2008). Hydrothermal vents and the origin of life. Nature Reviews Microbiology, 6(11):805–814.

Maury, C. P. J. (2015). Origin of life. primordial genetics: Information transfer in a prerna world based on self-replicating beta-sheet amyloid conformers. Journal of Theoretical Biology, 382:292–297.

Miller, S. L. and Urey, H. C. (1959). Organic compound synthesis on the primitive earth. Science, 130(3370):245.

Orgel, L. (2000). A simpler nucleic acid. Science, 290(5495):1306.

Orgel, L. E. (1968). Evolution of the genetic apparatus. Journal of Molecular Biology, 38(3):381–393.

Sakuma, Y. and Imai, M. (2015). From vesicles to protocells: the roles of amphiphilic molecules. Life (Basel, Switzerland), 5(1):651–675.

Schmitt, A. O. and Herzel, H. (1997). Estimating the entropy of dna sequences. Journal of Theoretical Biology, 188(3):369–377.

Segré, D., Ben-Eli, D., Deamer, D. W., and Lancet, D. (2001). The lipid world. Origins of life and evolution of the biosphere, 31(1):119–145.

Szostak, J. W., Bartel, D. P., and Luisi, P. L. (2001). Synthesizing life. Nature, 409(6818):387–390.

Vaidya, N., Manapat, M. L., Chen, I. A., Xulvi-Brunet, R., Hayden, E. J., and Lehman, N. (2012). Spontaneous network formation among cooperative rna replicators. Nature, 491(7422):72–77.

Walker, S. I. (2017). Origins of life: a problem for physics, a key issues review. 80(9):092601.

Wattis, J. A. D. and Coveney, P. V. (1999). The origin of the rna world: A kinetic model. The Journal of Physical Chemistry B, 103(21):4231–4250.

Wright, S. (1984). Evolution and the Genetics of Populations, Volume 1. The University of Chicago Press.

Wu, M. and Higgs, P. G. (2012). The origin of life is a spatially localized stochastic transition. Biology Direct, 7(1):42.

